# Genomic and metagenomic survey of microbial carbonic anhydrase genes reveals novel clades, high diversity, and biome-specificity

**DOI:** 10.1101/2025.01.09.631809

**Authors:** Mario E.E. Franco, Esther Singer, Simon Roux, Laura K. Meredith, Jana M. U’Ren

## Abstract

Carbonic anhydrase (CA) enzymes catalyze the interconversion of carbon dioxide and bicarbonate with an efficiency exceeded only by superoxide dismutase. CA enzymes have convergently evolved multiple times from phylogenetically distant organisms into structurally unrelated classes (α, β, γ, *δ*, *ζ*, *η*, θ, ι) with conserved physiological functions involved in photosynthesis, respiration, pH homeostasis, CO_2_ transport, and carbonyl sulfide hydrolysis that play central roles in medicine and the environment. Here, we leverage the recent surge in publicly available genomes and metagenomes to re-examine our understanding of the abundance, diversity, and phylogenetic relationships of the three major CA classes in Bacteria/Archaea and microbial Eukaryotes (Fungi, algae). We recovered a total of 57,218 α-, β-, and γ-CA sequences from 24,184 metagenomes and genomes, including the first detection of α-CA from an Archaeal species. CA sequences formed 3,859 protein clusters (1,188 with ≥ 3 sequences) that were taxonomically conserved at higher levels (i.e., Superkingdom, Phylum, Class). When viewed within a phylogenetic framework, the majority of subclades contained CAs representing multiple Superkingdoms, although numerous novel β-CA clades appear unique to Fungi. Queries of CA Hidden Markov models (HMMs) against all public meta-genome and -transcriptome datasets revealed that CA is a ubiquitous enzyme present in virtually all sampled environments. However, CA clusters that were taxonomically conserved also appeared environment-specific, generating high CA diversity. This work represents an important contribution to the evolution, diversity, and environmental distribution of an enzyme that is key for life and has broad environmental and industrial applications.

## Introduction

Organisms spanning all domains of life encode carbonic anhydrase (CA) enzymes that catalyze the hydration of carbon dioxide (CO_2_; CO_2_ + H_2_O ⇋ HCO_3_^−^ + H^+^) playing vital roles in carbon (C) fixation, respiration, CO_2_ sensing, pH regulation, growth, and reproduction [1–4]. Microbial eukaryotes such as red and green algae use CAs to concentrate CO_2_ around RubisCO, enabling them to photosynthesize in C-limited environments such as blooms [5]. CA activity impacts global biogeochemical cycles, by facilitating microbial roles in soil inorganic C formation and influencing isotopic signatures of CO_2_ [6–8] and sulfur (S)-containing gases in the atmosphere (e.g., carbonyl sulfide, OCS or COS) [9, 10]. Moreover, CA activity may fulfill organisms’ stoichiometric requirements (e.g., COS + H_2_O → CO_2_ + H_2_S) (e.g., [11]).

Unlike other key enzymes that share a single, common ancestor (e.g., bacterial multicomponent monooxygenase (BMM) family [12]), CA enzymes have evolved independently across diverse organisms. As such, CA represents one of the best examples of the convergent evolution of a common catalytic function in phylogenetically diverse organisms [13]. To date, at least eight CA classes have been described (α, β, γ, *δ*, *ζ*, *η*, θ, ι) [2, 5, 14–16]. Most organisms contain one or more CA classes [17] reflecting both intra- and inter-class redundancies [18]. Although maintaining proper levels of CO_2_, bicarbonate, and protons is critical for cellular processes [13, 19, 20], CA may be inactivated and/or lost completely for some organisms inhabiting high CO_2_ environments [21] or obligate, intracellular parasites [20]. In addition to convergent evolution, horizontal gene transfer (HGT) may influence an organism’s CA diversity. For example, organisms living in a marine hydrothermal vent appear to frequently exchange α-, β-, and γ-CA genes via HGT [22].

Understanding the diversity, distributions, and functions of different CA classes and isoforms is important for both medical and environmental applications. CA inhibitors, especially those that target specific CA classes or isoforms, are attractive drug targets for the treatment of human diseases (i.e., CA inhibitors are used as diuretics, anti-glaucoma, anti-convulsant, anti-obesity, anticancer agents/diagnostic [23–30]). As general CA inhibitors can cause side effects in humans, inhibitors that can specifically target bacteria or fungal CA isoforms are needed [17, 31, 32]. In addition, different CA classes/isoforms are associated with different ecosystem functions. In marine environments, phytoplankton upregulate specific CA (e.g., δ-, θ- and ι-CA associated with diatoms) as a C-concentrating mechanism to compete under low CO_2_ conditions [33–35]. In saline and freshwater wetlands, changes in the expression of β-A-CA with salinity and depth are associated with differences in dark CO_2_ fixation [36]. In soil, the bacterium *Thibacillus thioparus* and fungus *Trichoderma harzianum* encode a CA enzyme in the β-CA D clade (COSse), that can efficiently hydrolyze COS with relatively low CA activity for CO_2_ [37, 38].

A subset of specific CA classes and isoforms, encoded by diverse microbial organisms, may also drive patterns of atmospheric COS. COS directly impacts ecosystem sulfur availability, but also is a potential tracer for photosynthesis in terrestrial ecosystems due to the shared biochemical (CA) and physical (stomatal) pathway of COS and CO_2_ in leaves [10, 39]. As demonstrated by inhibitors (e.g., [40]), soil microbes exhibit significant CA activity and the ability to degrade COS [41–43]. Soil COS-degrading microbes tend to encode for β-D-CA [44, 45], which appear to be highly expressed in soil metatranscriptomes [42]. Fungal CA appears to be particularly important for soil COS uptake [41, 42, 46], which may explain reductions in soil COS uptake under N fertilization [47] via shifts in fungal diversity [48] and high COS uptake rates in fungal-dominated leaf litter [49]. Similarly, CA activity, likely of α-CA of certain algal and bacterial taxa [41, 42], influences the atmospheric budgets of δ^18^O in CO_2_ and H_2_O are therefore important to consider during the use of these tracers of large-scale patterns in the C cycle and precipitation [41, 42, 50–56].

Despite the ubiquity and importance of CA, few metagenomic and metatranscriptomic studies have specifically examined the diversity and abundance of CA in the environment. As such, the overall abundance, diversity, and distribution of CA across diverse organisms (representing both cultured and uncultivated) and terrestrial and marine ecosystems remains largely unknown. Here, we use large-scale public genomic data to perform a systematic assessment of CA spanning both prokaryotic and eukaryotic microbes. By inferring patterns of CA across genomes and environments, we provide a framework to better understand CA abundance and taxonomic conservation across the tree of life that links CA with its key physiological, ecological, and environmental roles (e.g., as volatile traits [57]) that influence atmospheric C and S budgets.

## Materials and Methods

### Database Mining, Sequence Clustering, and HMM Building

Based on functional annotation of α-, β- and γ- CA derived from Clusters of Orthologous Groups (COGs) [58] and Protein Families (Pfam), we downloaded putative α-, β- and γ- CA protein sequences from Integrated Microbial Genomes (IMG) [59] in May 2021 using three COGs IDs COG3338, COG0288, and COG0663; and from MycoCosm [60] and PhycoCosm [61] using three Pfam [62] IDs pf00194, pf00484, and pf00132. These COGs and Pfam IDs were chosen based on previous work by Meredith et al. [42] that recovered assembled α-, β- and γ-CA genes from soil metatranscriptomes using HMMs built with the above COG IDs. Since MycoCosm and PhycoCosm lack COG annotations, we use the Pfam IDs associated with the above COG IDs in the JGI MycoCosm [60] and PhycoCosm databases (Table S1).

CA sequences were retrieved from 24,184 genomes or metagenome-assembled genomes (MAGs) representing 85 bacterial, 12 archaeal, 6 algal, and 9 fungal phyla. After length filtering (see Supplementary Methods) CA sequences were clustered using MMseqs2 [63] at 50% amino acid (aa) similarity. Following clustering, we constructed multiple sequence alignments (MSAs) with MAFFT v7.453 (*--auto*) [64] and used HMMER 3.3.2 [http://hmmer.org/] to generate a Hidden Markov Model (HMM) for CA clusters with at least three sequences. Taxonomic information for each sequence was used to determine the lowest common ancestor (LCA) of each CA cluster (i.e., the lowest taxonomic level shared by all members of the cluster) [65].

### Multiple Sequence Alignments and Phylogenetic Analyses of CA

We combined the representative sequence of each CA cluster (provided by Mmseqs2 [63]) with reference CA sequences derived from previous studies [2, 37] and records in UniProt/SwissProt with the highest level of annotation evidence [66] (hereafter, UniProt/SwissProt reference sequences). Representative sequences of each CA cluster were aligned to the reference alignment for each CA clade using the *--addfragments* option with MAFFT v7.453 [64]. MSAs were then trimmed using trimAl version 2 [67] with the *--gappyout* option, followed by visual inspection and alignment adjustment in Geneious version 2021.02.02 [68]. We removed 257 (220 γ-CA) sequences representing CA clusters that appeared fragmented (<75% of alignment length) or lacked the characteristic CA metal binding sites [2, 69–71]. The remaining sequences were re-aligned (237 α- CA, 521 β-CA, and 284 γ-CA sequences) as described above and phylogenetic analyses were performed with IQ-TREE multicore version 1.6.12 [72] using the best-fitting model of sequence evolution selected by ModelFinder [73] and 1000 ultrafast bootstrap (UFBoot) replicates [74]. Nodes with less than 95% UFBoot support were collapsed [75, 76]. CA phylogenetic diversity (PD) (i.e., sum of all the branch lengths) was computed for the entire tree and for only fungal sequences using the R package *caper* version 1.0.1 [77], with randomizations as described by ref [78].

### Enviromental distribution of CA

We queried 1,188 CA HMMs against 3,688 published metagenome (metaG) and metatranscriptome (metaT) datasets from JGI IMG [59]. Database hits to CA HMMs were filtered based on a score ≥ 50. The highest-scoring open reading frame (ORF) from a metaG/metaT was assigned to a single CA HMM. CA hits were used as a proxy for abundance, scaled by study proportion per ecosystem (P) and taxonomic group abundance (T). Scaled values were log-transformed (LOG((P/T)+0.00001)) and visualized using heatmaps with the R package *pheatmap* v 1.0.12 [79]. For simplicity, we collapsed the detailed ecosystem categories into seven categories: engineered environment, air, aquatic/marine, terrestrial: other, terrestrial soil, plant-associated, and other host-associated (e.g., insect gut).

## Results

### Variation in CA distributions, identities, and copy number

We compiled 57,218 putative α-, β-, and γ-CA sequences from the JGI databases (Table S3, Additional file 1). After filtering potentially misannotated sequences, 54,631 sequences remained (Table 1). Over 70% of genomes/MAGs contained CA proteins. Most of the genomes lacking CA derived from uncultured Bacteria and Archaea (i.e., MAGs) (Fig. 1, Fig. S1) Seventeen fungal genomes without detectable CAs represented four different phyla (Fig. S1, Table S4). *Guillardia theta* was the only algal genome without detectable CA (Fig. 1, Table S4).

**Figure 1.**
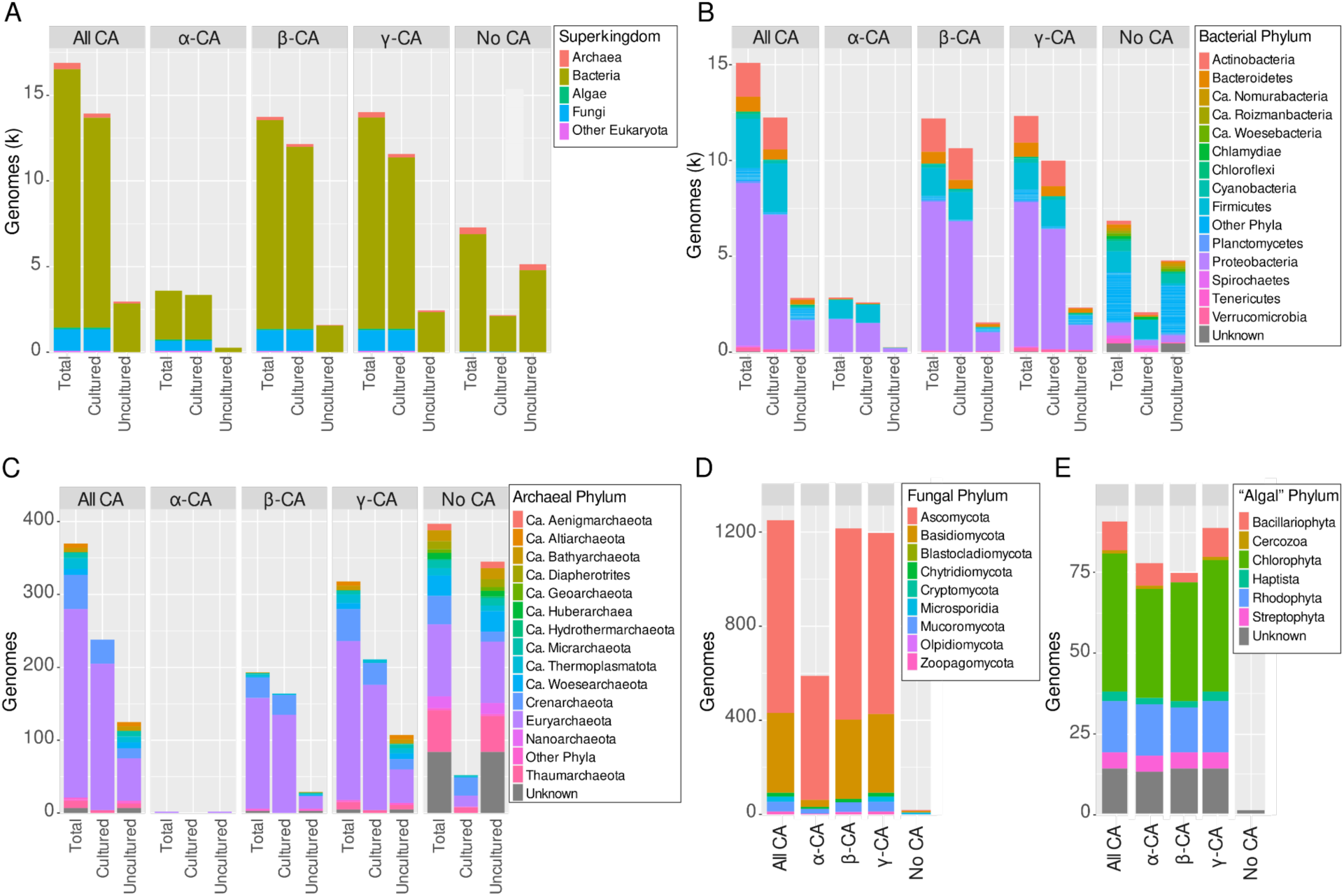
Number of genomes containing CA genes and their taxonomic affiliation by (**A**) Superkingdom and phylum for (**B**) Bacteria; (**C**) Archaea, (**D**) Fungal, and (**E**) algae. In panel B, less abundant bacterial phyla were grouped together as “Other Phyla” for visualization purposes. For Bacteria and Archaea, the number of CA for cultured vs. uncultured organisms (i.e., MAGs) is also shown. Fungal and algal genomes from Mycocosm and PhycoCosm are only represented culturable organisms. Genomes of “algae” in PhycoCosm represent green algae (Chlorophyta), red algae (Rhodophyta), as well as a diverse assemblage of unicellular eukaryotes, including diatoms (Bacillariophyta), amoeboids (Cercozoa), haptophytes (Haptista). Also see Table S3.

**Table 1.**
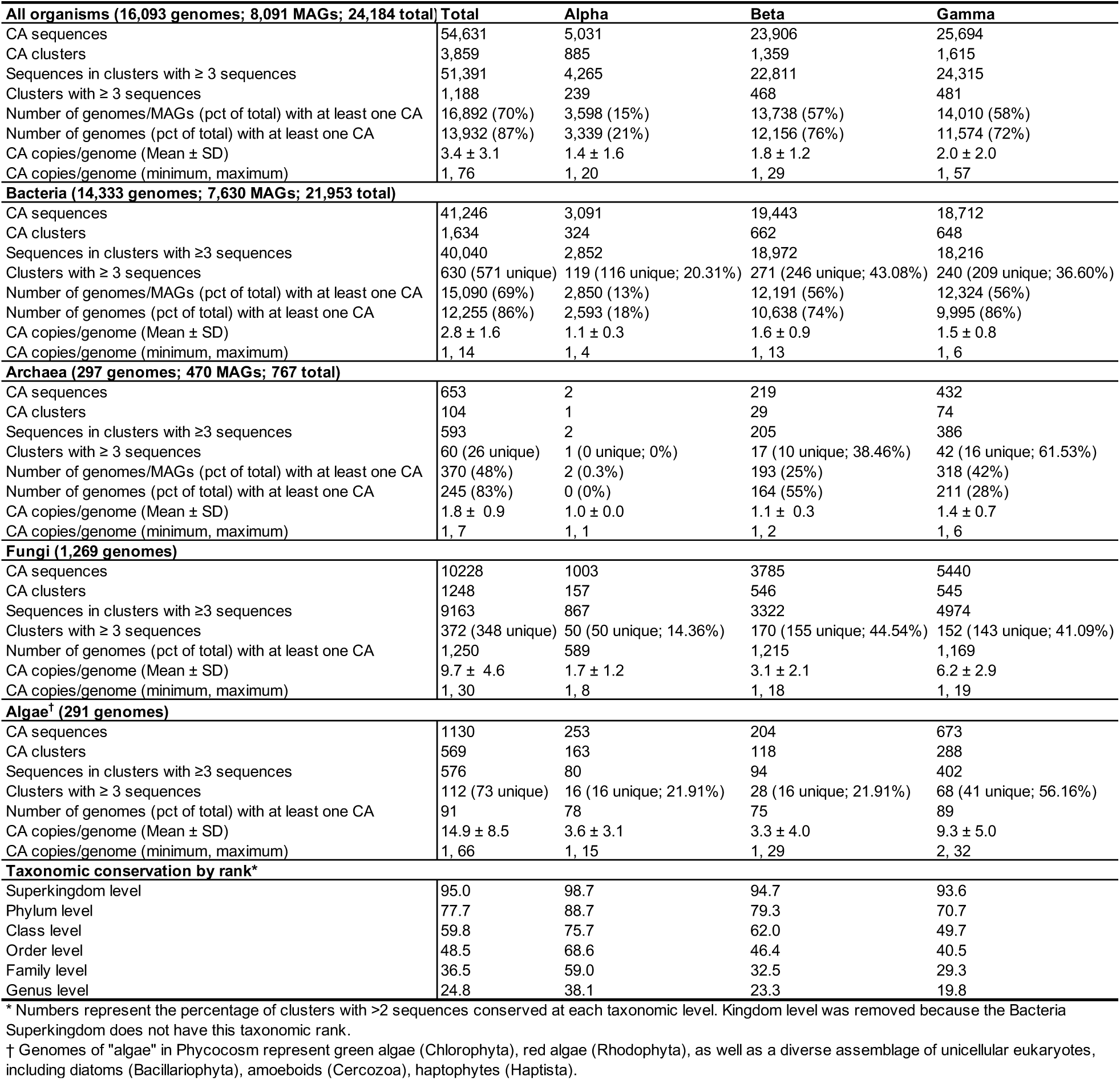
Summary of CA sequences retrieved, degree of sequence diversity (clusters), copy number variation, and taxonomic conservation at various ranks by CA class.

CA presence varied by class and taxonomy, with 98.5% of fungal genomes containing CAs, compared to 68.7% of bacterial, 48.2% of archaeal, and 31.3% of algal genomes. However, the percentage of bacterial and archaeal genomes with at least one CA increased to 85.5% and 82.5%, respectively, when restricted to genomes from cultured organisms (Table 1, Table S3b). α-CAs were found in the smallest fraction of genomes (14.9%). Although α-CAs have not been previously detected in Archaea [13, 80], we detected one putative α-CA sequence in two MAGs of *Archaeoglobus* [81] (Fig. 1a, Table S3, Additional file 1). A higher fraction of genomes contained β- or γ-CAs, and both classes were found in similar percentages of genomes within different taxonomic groups (Table 1).

CA-encoding genomes averaged 3.4 copies/genome, with algae having the highest (14.9 ± 8.5), followed by fungi (9.7 ± 4.6), and lower numbers in bacteria (2.8 ± 1.6) and archaea (1.8 ± 0.9) (Table 1, Table S5, Fig. 2). The basidiomycete fungus *Fibulorhizoctonia psychrophila* CBS 109695 (MycoCosm portal: Fibsp1) had the most copies (7 α-, 13 β-, and 10 γ-CAs) (Table S5, Additional file 1). In genomes with multiple CA classes, a combination of both β- and γ-CA was the most common (Fig. 3). Algal genomes had the highest frequency of all three CA classes, followed by Bacteria and Fungi. In contrast, no archaeal genomes contained all three CA classes and instead were more likely to contain only γ-CA (Fig. 3). No Fungal or Algal genomes contained only α-CAs.

**Figure 2.**
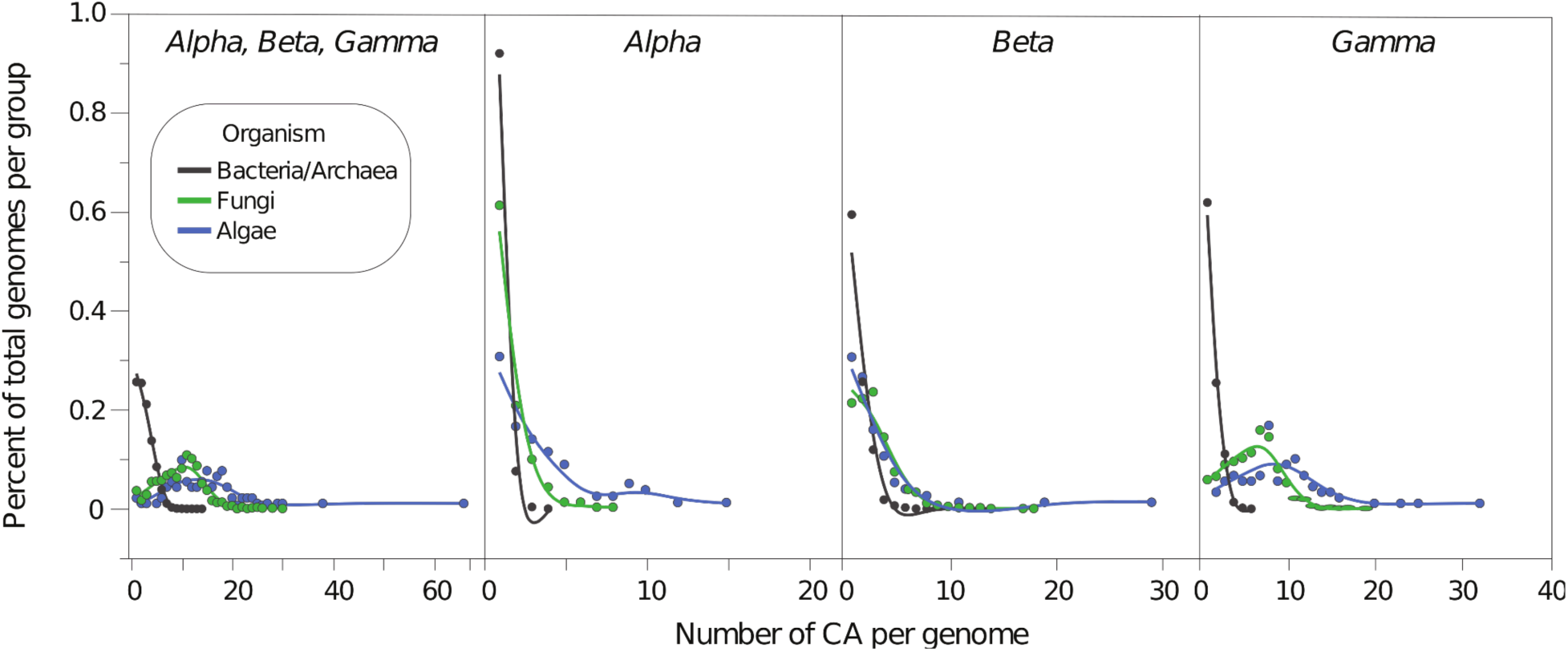
CA copy number variation among different microbial taxonomic groups. The percent of total genomes per taxonomic group with different CA copy number for all CA classes combined and for each CA class individually. Points and lines are colored by organismal taxonomy (see legend).

**Figure 3.**
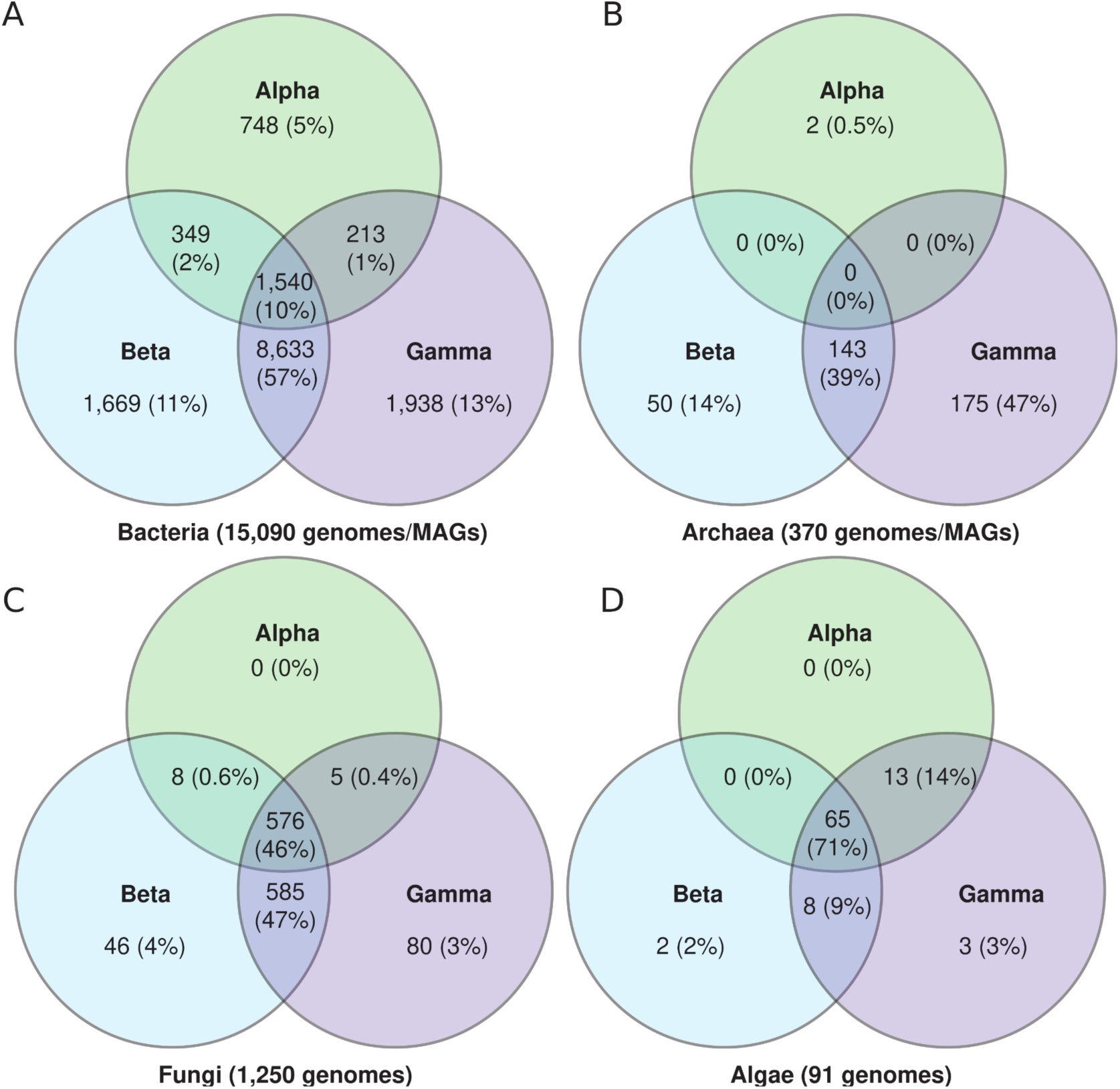
Distribution of different CA classes within microbial genomes. Venn diagram of the number (and percentage of total CA) of α-, β- and γ-CA within genomes of Bacteria, Archaea, Fungi, and algae.

### Diversity and taxonomic conservation of microbial α-, β-, and γ-Cas

To assess CA diversity among prokaryotic and eukaryotic microorganisms we clustered all 54,631 CA sequences at 50% amino acid identity, which resulted in 3,859 CA clusters, mostly singletons (Table 1, Fig. 1b, Table S6, Additional file 2). Although the number of bacterial genomes/MAGs exceeded eukaryotic genomes 14-fold, the number of CA sequences from Bacteria was only 3-fold greater than eukaryotic genomes (Table 1). Both Superkingdoms had similar cluster counts after excluding clusters with fewer than three sequences, whereas the smaller number of Archaeal genomes resulted in correspondingly fewer clusters (Table 1). CA clusters with ≥3 sequences were highly conserved at the Superkingdom level, although conservation declined at lower taxonomic ranks (Table 1; also see Table S7, Additional file 3). α-CA clusters were the most taxonomically conserved at all ranks (Table 1). Uncertain rank (i.e., *incertae sedis*) or missing taxonomic information had a negligible impact on LCA assignment (see Additional file 3).

### CA phylogenetic relationships, diversity, and taxonomic conservation

The maximum likelihood phylogenetic analyses of α-CA and β-CA resulted in numerous well-supported clades (UFBoot values >95%), with tree topologies and clades consistent with prior studies (Fig. 4, Figs. S4-S5, Figs. S7-S8, Additional file 4). However, relationships among most γ-CA clusters could not be resolved with high statistical support (Fig. 4, Fig. S6, Fig. S9, Additional file 4). Our analyses reveal a large degree of novel CA diversity within each previously described CA class, and also revealed several novel CA clades (Fig. 4). Here, we delineated five new clades of α-CA (α A-E) (Fig. 4b). Additionally, we described a new β-CA clade (β-E) and three subclades of β-D (β-D1, β-D2, β-D3), with fungal CAs found in β-A and β-D clades (Fig. 4a). Fungal CAs were found in two β-CA clades (β-A and β-D). The β-D2 clade contains previously identified sequences for bacterial and fungal COS-hydrolyzing COSase enzymes [2, 37] and fungal Cas3 enzymes (also in β-D3) [82] (Fig. 4a). Although the phylogeny of γ-CA lacked strong bootstrap support, the resulting maximum likelihood topology supports at least four major clades of γ-CA (Fig. 4c). α-CA showed the lowest phylogenetic diversity (PD), whereas β-CA had the highest, with fungal sequences significantly increasing PD across all classes (P < 0.001) (Fig. 4d).

**Figure 4.**
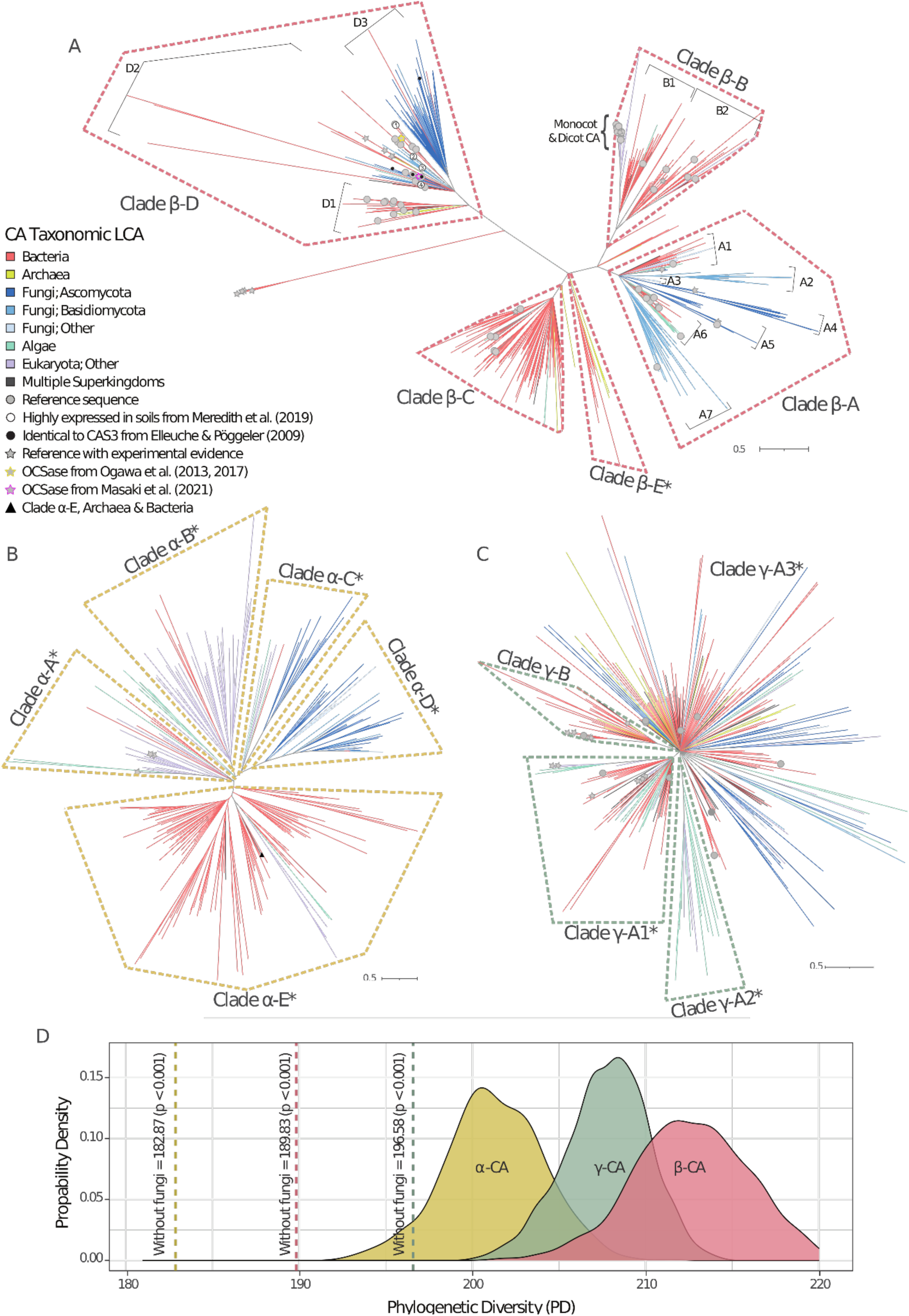
Phylogenetic analysis of α-, β-, and γ-CA reveals novel diversity. Phylogenies of (**a**) β-, (**b**) α-, and (**c**) γ-CAs based on ML analysis. Node support represents 1000 ultrafast bootstrap (UFBoot) replicates. Nodes with less than 95% UFBoot support were collapsed (see Supplemental Figs S4-S6 for phylogenies without collapsed branches). Newly described subclades of each CA class are labeled with asterisks (*) (e.g., clade β-E*). Branches are colored according to Superkingdom or Phylum based on the taxonomic LCA of each CA cluster. The scale bar represents the number of substitutions per site. Different symbols indicate significant CA reference sequences from previous studies of CA (see legend). There were three α-, 85 β-, and 22 γ-CA reference sequences. All three α-CA reference sequences were from UniProt/SwissProt. β-CA reference sequences included 27 CA proteins from UniProt/SwissProt, 32 from Smith et. al. [2], and 41 from Masaki et al. [37]. γ-CA reference sequences were comprised of 11 UniProt/SwissProt sequences and 14 sequences from Smith et al. [2]. A small number of reference sequences were redundant across databases/studies (i.e., sequences used previously (Table S2). Overall, the placement of all β-A and β-C CA reference sequences was in agreement with previous analyses. One exception was sequence ZP_00310115 that our analyses placed in the β-C clade, but was previously placed in the β-A by Masaki et al. (2021) (Fig. 4, Figs. S5, S8). (**d**) Probability distribution of the phylogenetic diversity of each CA class. The discontinuous line indicates the level of PD obtained when fungal sequences are excluded from analyses. P-values were calculated using 1000 random subsets of non-fungal clusters in such a way that the sample size equals the total number of non-fungal clusters.

### Abundance and distribution of CA clusters in the environment

We queried 1,188 CA HMMs against 3,688 published metagenome (metaG) and metatranscriptome (metaT) datasets from JGI IMG [59]. The CA HMMs had hits in 99.5% of all published studies on IMG. 85% of CA clusters had at least one hit across all 3,672 metagenomic and metatranscriptomic datasets. Most clusters without hits were assigned to a Eukaryotic LCA, with β-CAs being the most common (Fig. S10). We found no observable relationship between CA cluster size and the number of hits/cluster (Fig.S10). For α- and β-CAs, clusters that were not conserved at the Superkingdom level occurred in a significantly higher mean percentage of datasets and had a greater mean number of hits to the dataset compared to CAs that were assigned to a single Superkingdom (P<0.0001) (Fig. S12).

Aquatic and soil ecosystems had the highest CA hits (Fig. 5a, Table S8). γ-A CAs were the most abundant CA class across all environments (except in host-associated metatranscriptomic datasets) (Fig. 5a, Fig. S13). In contrast, α-CAs were the least abundant overall and absent in certain cases such as host-associated, engineered, or freshwater aquatic environments (Fig. 5a, Fig. S13). β-CA clades A-D occurred in similar frequencies across most environments, except for freshwater aquatic and engineered environments that only contained β-D or β-A, respectively (Fig. 5a). In contrast, β-E CA had few hits to any environment.

**Figure 5.**
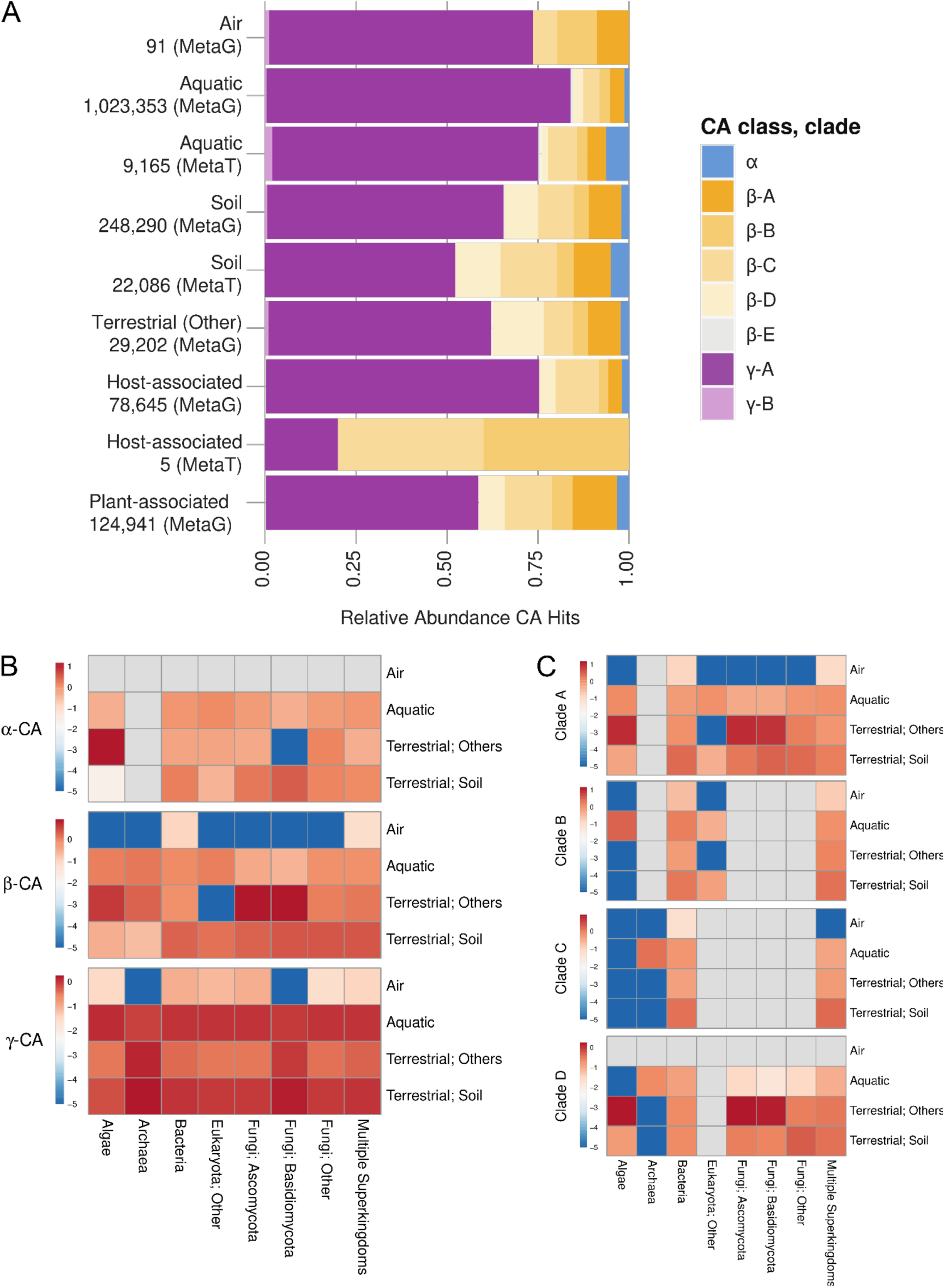
Relative abundance of different CA classes and clades by environment. Data are based on α-, β- and γ-CA HMM hits to the metagenomic/metatranscriptomic databases at IMG. (**A**) Stacked bar graph of the relative abundance of each CA clade by environment. Numbers represent the number of hits per environment for either database. Metatranscriptomic information is only available for aquatic, soil, and host-associated environments. Clade information follows Fig. 4. Host- associated data are primarily from studies of arthropod gut microbiomes. Heatmaps showing the relative abundance of CA across four major environments by taxonomic group (determined by LCA) for (**B**) α-, β- and γ-CA clusters and (**C**) four clades of β-CA clusters. The number of hits was scaled by the proportion of studies for the respective ecosystem and the abundance of each taxonomic group, and log-transformed to account The data were normalized to account for potential bias due different numbers of studies per ecosystem (i.e., greater number of studies from marine and soil environments) and towards bacteria/archaea whose smaller genomes may be more readily sequenced than eukaryotic organisms.

Overall, terrestrial environments and soil had a higher number of fungal CA hits, whereas the number of bacterial/archaeal CA hits was consistent across environments (except for air) (Fig. 6; set size). Bacterial/Archaeal CA from each environment was dominated by CA with sequences from multiple phyla and Proteobacterial CA (Fig. 6a), whereas Fungal CA from each environment was dominated by Ascomycota CA (Fig. 6b). Although relative abundances of CA assigned to different bacterial/archaeal and fungal phyla were similar among ecosystems (i.e., see set size), the majority of CA clusters appear to be specific to either one or two environmental categories (Fig. 6, Fig. S14, Table S9). There was no clear association between CA clade and environmental distribution (i.e., all CA clades were found in most environments) (Fig. 6).

**Figure 6.**
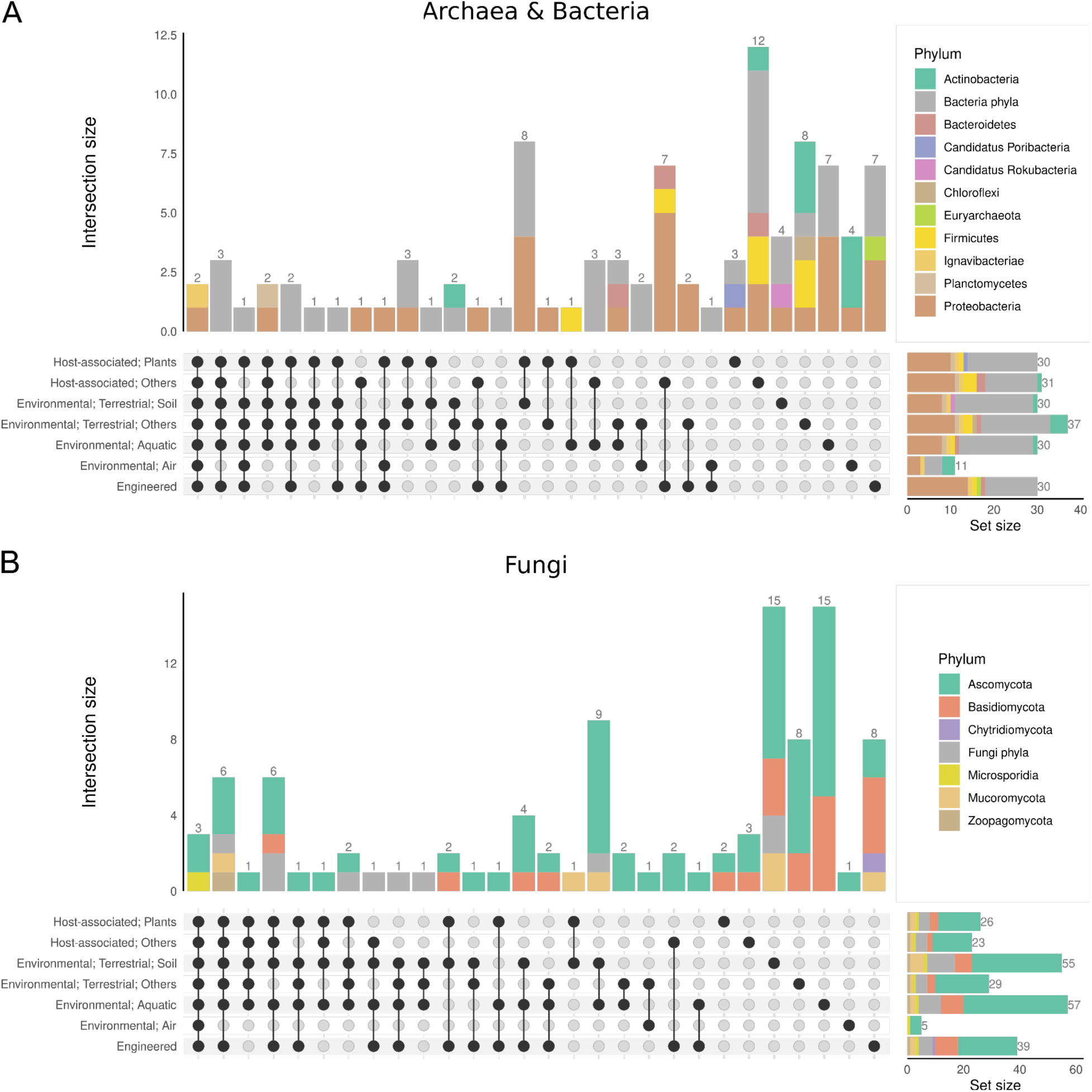
Specificity of CA HMMs to different environmental categories. UpSet plot displaying the intersection of different ecosystems for the ten most abundant HMMs per environment for (**A**) Archaea and Bacteria and (**B**) Fungi. Each vertical stacked bar shows the Phylum-level composition and number of CA clusters with that combination of environments (i.e., intersection). Horizontal stacked bars show the Phylum-level composition and number of CA hits (i.e., set size) for each environment. Phylum designation is based on LCA taxonomic assignment. The most abundant HMMs are listed in the Table S9.

## Discussion

### New insights into expanded α-CA diversity in Gram-positive bacteria and the Archaea superkingdom

Phylogenetic studies suggested γ-CA evolved first (arising ca. 4.2 billion ago [2]) followed by the independent evolution of β-CA (no age estimate due to accelerated divergence [2, 83]) and then α-CA (arising ca. 0.5-0.6 billion years ago [71]). Among the previous observations supporting this hypothesis was the resolution of a monophyletic clade containing all the α-CA sequences nested between β-CA sequences of Gram-negative Bacteria, coupled with the fact that α-CA had previously only been found in Gram-negative Bacteria [4, 80, 84] until it was recently reported in the Gram-positive *Lactobacillus rhamnosus* GG [85]. We found that ca. 25% of Gram-positive bacterial genomes contained α-CA (Additional file 1). Gram-positive bacteria genomes with α-CA included 63 different genera (e.g., *Listeria monocytogenes*, *Lacticaseibacillus*, *Oenococcus oeni*, *Streptococcus*, *Enterococcus*, *Streptomyces*, and *Bacillus*). In addition, although archaeal genomes primarily contained only γ-CAs, we detected α-CA genes in two MAGs of the archaeal genus *Archaeoglobus* obtained from fluids collected deep within the Juan de Fuca Ridge subseafloor [81] (Table 1, Table S3b, Additional file 1)—a genus known to contain thermophilic sulfate reducers [86, 87]. To the best of our knowledge, α-CA has never been reported in Archaea [13, 80]. Noteworthy, eleven other *Archaeoglobus* genomes lacked α-CA genes. Thus, while our analyses are consistent with *Archaeoglobus* acquiring α-CAs via horizontal gene transfer (HGT) from Bacteria, additional analyses are necessary to confirm this finding. While we find α-CA to be the most conserved and least phylogenetically diverse of these CA, our large-scale genomic analysis also uncovers a wider distribution of α-CA in prokaryotes than previously appreciated.

### Organisms exhibit diverse CA genome content

Reflecting their larger and more complex genome architecture [88–90], fungal and algal genomes contained 3.8 to 5.9 times more CA copies/genome on average than bacterial and archaeal genomes, respectively (Table 1, Table S5). Although a relatively high proportion of bacterial and archaeal genomes have multiple copies of β- and γ-CAs, almost half of all fungal genomes (46%) contained all three classes of CA, as well as duplications within each class (S2d Table). In eukaryotic genomes, there are on average 2X more copies of γ-CA compared to β-CA per genome, while α-CA is usually not duplicated within a genome. However, the differences are more subtle in prokaryotic genomes, where the β-CA and γ-CA classes are highly abundant in Bacteria and Archaea, respectively (Table 1, Table S5).

Although widespread, some microbes appear to completely lack CA. One such example is the syntrophic bacterium *Symbiobacterium thermophilum,* which grows on CO_2_ generated from other bacteria [91]. The fact that closely related *Clostridia* (Firmicutes) contain CA suggests that *S. thermophilum* has lost the enzyme as an adaptation to a high CO_2_ environment. Few other CA-deficient organisms have been reported [2, 17], although an analysis of published Proteobacteria genomes found 39 strains lacking CA [21]. Similar to *S. thermophilum*, it is hypothesized that loss of CA in these organisms is associated with a lack of selection for CA due to living in high-CO_2_ environments [21].

Dominant culturing approaches under relatively low (ambient) CO_2_ conditions may select against CA-deficient organisms that require high CO_2_ for growth and contribute to these patterns [21]. Host-associated organisms may be more likely to live in high CO_2_ environments, facilitating continued growth with the loss or inactivation of CA (e.g., [20]). CA inhibitors are a growing area of interest for anti-bacterial and anti-fungal treatments [31, 92], with the idea that they are central to the function of these pathogens [30]. However, we find evidence for a widespread absence of CA, suggesting that some microbes not only continue to survive (e.g., fungal CA knockouts survive under low CO_2_ conditions but with strongly reduced fitness/growth rates [93]) but also thrive even when lacking CA. Genomes lacking genes annotated as CA included phylogenetically diverse taxa (Fig. 1, Fig. S1), consistent with repeated loss of CA across lineages. Many single-cell genomes and MAGs correspond to poorly understood taxa, so the full suite of environmental factors contributing to CA loss is unclear.

Known alternate CA classes do not fully explain the lack of CA in these organisms. Some apparent CA-deficient genomes identified here may encode for one or more of the other five described CA classes (i.e., δ-, ζ-, η-, θ- and ι-CA), however the phylogenetic distribution of most of these classes is thought to be highly restricted to mainly diatoms and protozoa, and BLAST searches for δ-, ζ-, η-, and θ-CA resulted in <2% of putative CA recovered for α-, β-, or γ-CA (Table 1, Table S1) and thus cannot explain the large proportion of genomes observed without a CA. The ι-CA appears more broadly phylogenetically distributed compared to δ-, ζ-, η-, and θ-CA [16, 34], but likewise recovered <1% of the putative CA from the IMG database via a BLAST search and is unlikely to explain the widespread lack of CA observed. Nevertheless, the ongoing discovery of new CA, including the ι-CA suggests that novel CA and their diverse roles in organisms could still be undiscovered.

### Phylogenetic conservation of CA is strongest at the sub-class level

CA sequences (even within a class) are poorly conserved compared to other enzymes [83, 94, 95], as shown by the high number of CA clusters at 30% and 50% aa similarity (Table 1, Fig. 2; see also Table S6, Additional file 2). Phylogenetic analyses revealed many unsupported nodes and polytomies, reflecting high sequence divergence and the challenge for short protein sequences to represent deep evolutionary history (Fig. 4; also see Fig. S4-S9).

Overall, γ-CA phylogenetic relationships were the most difficult to reconstruct. Previous research suggests that the majority of extant γ-CAs arose from multiple simultaneous divergence events that took place relatively close to the last common ancestor (LCA) of all γ-CAs [2]. This may indicate an ancient diversification of γ-CA potentially associated with different microbial clades and ecological niches. Interestingly, the CA lineages diverging from the largest polytomy in the phylogeny were found to have LCA representing all major taxonomic groups (Eukarya, Bacteria, Archaea). Since γ-CA is hard to identify based on Pfam and COG information due to diverse functions, we removed sequences misaligned with validated γ-CA metal-binding sites, reducing the sequences used for γ-CA phylogeny (Table S2c). There also could be more flexibility in CA functionality than currently appreciated. For example, despite the classic understanding of CA as strict metalloenzymes, CA from the most recently described class (ι-CA) retains activity even in the absence of a metal cofactor [96]. Future research is needed to functionally validate the CA enzymatic activity of highly divergent γ-CA would assist in the development of better methods to identify γ-CA using sequence-based methods.

Unlike γ-CA, phylogenetics of α-CA and β-CA had stronger statistical support and well-supported clades (Fig. 4; see also Fig. S4-Fig. S9). Phylogenetic analyses also showed more genetic changes in β-CA compared to other clades, possibly due to the more recent divergence of the α-class [2, 71]. Previous reconstructions indicate frequent duplications and horizontal gene transfers in β-CA, likely due to its modularity [71, 83]. By including LCA taxonomic information for the CA sequences, we found that the majority of well-supported clades had LCAs representing a single Supergroup. For example, numerous Fungal- or Bacterial-specific clades are evident in both the α- and β-CA phylogenies (Fig. 4). The β-CA phylogeny recovered most of the clades introduced by Smith et. al. [2] and at least two additional clades. Our analyses suggested some α- and β-CA clusters align with microbial taxonomy, enabling CA diversity estimation via taxonomic profiles [97].

### Fungi and algae are important reservoirs of CA diversity

Genomic data integration revealed an outsized reservoir of CA diversity in eukaryotic life on a per-genome basis. Fungi had the most CA copies per genome and the inclusion of fungal genomes significantly increased known CA diversity, and were dominant members of new CA subclades. Fungal CA are involved in growth and sexual development [93] and CO_2_ sensing, which is a critical capability for pathogenic fungi that experience wide-ranging CO_2_ concentrations [98] and fungi modify expression of virulence traits accordingly [31, 99]. CA duplication may reflect their diverse roles and cellular locations [82], with varied catalytic activity [93, 100]. Fungal CA abundance rivals that of bacteria for some CA types and environments (Fig. 5), yet their role in ecosystem processes has received little investigation thus far (but see [42]).

Here, we confirm the widespread presence of α- and β-CA in Fungi. While the previous focus in Fungi was specific β-CA isoforms [82, 92, 101, 102], including those similar to plants (Cas1 and Cas2) but also to Bacteria (Cas3) [82], our analyses show that fungal CAs are part of a much bigger picture (Fig. 4) (see also [11]). We also infer abundant γ-CA in Fungi, which has received little attention despite being widespread and also duplicated in Fungal genomes (Figs. 1-2) [31, 82, 101]. However, experimental evidence is lacking to demonstrate the CA function of Fungal γ-CAs that are present only in the largely unresolved γ-A3-CA clade. The lack of resolution in this clade may arise from a history of rapid divergence, and γ-CA evolutionary history may be best reconstructed using protein structures. Understanding the distribution of CA across Fungi clades will help inform ongoing efforts to develop CA inhibitors as drug targets for fungal pathogens [31].

Finally, our analysis also demonstrated that algae contain nearly equal numbers of α-, β-, and γ-CA at a high level of duplication, and these algal CA (alongside other eukaryotic CA) may form novel subclades of CA diversity (e.g., γ-A2). Despite the potential to underestimate contributions of Eukaryota using environmental shotgun sequencing, we still find high expression of algal α-, β-, and γ-CA in some environments that will complement activity of their more specialized CA (e.g., δ-, θ-CA). Holistically representing eukaryotic CA represents an exciting frontier for understanding CA function and ecology in the environment.

### CAs are widely distributed across environments

Overall, we found that environmental CA gene content (metaG) and gene expression (metaT) were typically dominated by γ-CA followed by a sizable fraction of diverse β-CA and low to negligible levels of α-CA (Fig. S13). Previous analysis of 10 metatranscriptomes found a higher proportion of β-CA than γ-CA and relatively small fraction of α-CA in soils [42]. One explanation for this difference is methodological, as here we describe the diversity of CA functional potential in these environments rather than quantifying transcript abundance [42]. In addition, the abundance of γ-CA hits in metaT/G datasets compared to previous metaT of soils may be the result of including clusters with fewer sequences (i.e., 481 clusters with ≥3 sequences here vs. 81 clusters with ≥50 sequences from ref [42]). However, we found similar numbers of small clusters for all CA classes (Supplementary Information). Additional work is needed to experimentally verify the function of γ-A3-CA to refine inferences from ‘omics data.

Comparing the relative abundance of each CA in terms of gene content (metaG) versus expression (metaT) reveals that β-CA tended to be relatively more expressed in aquatic, soil, and host-associated data sets and α-CA expression in aquatic and soil environments but not host-associated, while γ-CA were always expressed at a relatively lower proportion compared to the available gene content. This suggests that while γ-CAs are the most abundant CA in genomes and environmental gene content, organisms may invest more heavily in β-CA and α-CA enzymes, implying outsized functional roles for these CA classes in the environment. Generally, we find lower β-A-CA content and expression in saline vs fresh aquatic systems (Fig. S13), consistent with a recent paired metaG/metaT study in wetlands exploring their role in dark CO_2_ fixation [36], and β-A-CA levels tended to be highest in terrestrial, plant-associated, and air environments (Fig. 5), which may be interesting environments to explore chemosynthetic C-fixation. We find that algae contain similar proportions of α-, β-, and γ-CA but are somewhat patchy in their relative abundance in different environments (Fig. 5); a deeper analysis of the regulation patterns of algal CA for these three cosmopolitan CA versus more conserved algal moieties (i.e., δ-, ζ-, θ- and ι-CA) could give insight into drivers of CA evolution and diversification. We queried each of the CA HMMs against the IMG public metaG/metaT database to investigate the relationship between the taxonomic affiliation of CAs and their environmental distribution. Lastly, although HMMs with the most hits represented taxonomically diverse CAs, they were typically specific to one or two ecosystems, which suggests that particular CAs may be shared across different microbial lineages due to environmental selection for particular CA functions.

### A refined picture of carbonic anhydrase in soil and its influence on the global atmosphere

Our approach highlights a convergence in the current understanding of the specific genes and organisms driving the global soil sink for atmospheric COS. We expand previous genome database-mining efforts [11, 42], leveraging databases with extensive prokaryotic, fungal, and algal genomes and implementing a new protein clustering approach [42] to contextualize COS degradation among the diversity of CA. Specifically, we localize known COS activity to the β-D2-CA subclade (Fig. 4), which contained both bacterial and fungal experimentally validated COSase sequences [37, 38], the primary CA clusters previously found to be expressed in soils [42], sequences identical to the cab-type β-CAS3 CA described in fungi [82], as well as experimentally validated reference sequences. Membership in the nested β-D2-CA clade included taxa previously associated with CA activity for COS through culture-independent (Ascomycota, Basidiomycota, Actinobacteria, and Proteobacteria [42]) and culture-dependent methods [37, 38, 44, 45, 103–107]. In contrast, the β-D3-CA nested clade was formed almost entirely of fungal sequences especially within the Ascomycota and there was only one bacterial CA cluster. An amino acid tree of COSase-like genes derived from the JGI MycoCosm by [11] recovered a similar Ascomycota-dominated clade despite different methods (i.e., recovery using blastp E-value 10^−25^ with THIF08 COSase as a query). Here, we recover multi-kingdom microbial CA diversity and reveal a more balanced role for prokaryotes among the β-D diversity, including a β-D2 nested clade containing nearly equal representation of prokaryotes and fungi, which differs from the concept of a Basidiomycota-dominated β-D2 clade [11], and we also describe a third β-D1 clade primarily consisting of prokaryotes. Our analyses of IMG metaG/T database provide additional evidence for the presence and expression of microbial β-D-CAs in soils and suggests that fungi in the Mucoromycota also may contribute to COS fluxes in soil (Figs. S14d, S15d, S16d). Here, we observe a relatively higher proportion of β-A-CA and β-C-CA in soil compared to the previous metaT analysis in [42]. β-D-CA, the most divergent CA [2, 83], includes neofunctionalization (e.g., to a COS hydrolase and carbon disulfide hydrolase) potentially used by fungi to obtain atmospheric sulfur when COS levels were higher in the atmosphere [11]. This new genomic resource developed here (e.g., HMMs) will enable future studies to test specific hypotheses on the role of the β-D2-CA nested clade in COS consumption in soils.

## Acknowledgments

This work was supported by funding from NSF AGS Atmospheric and Geospace Science Atmospheric Chemistry Program (Award number 1933280) to L.K.M. and J.M.U., as well as the University of Arizona Technology and Research Initiative Fund (TRIF) funds to J.M.U. Work conducted by E.S. and S.R. at the U.S. Department of Energy Joint Genome Institute (https://ror.org/04xm1d337), a DOE Office of Science User Facility, was supported by the Office of Science of the U.S. Department of Energy operated under Contract No. DE-AC02-05CH11231.

## Data Availability Statement

Data files used in analyses are available in FigShare repository (DOI: 10.6084/m9.figshare.28156742).

