## Supplemental Materials for "Genomic and metagenomic survey of microbial carbonic anhydrase genes reveals novel clades, high diversity, and biome-specificity"

### ***METHODS***

***Carbonic Anhydrase Database Mining, Sequence Clustering, and Hidden Markov Model Building.*** Reference α-, β-, and γ-CA sequences range from 151 to 687 amino acids (aa), while putative CA sequences range from 30 to 5,857 aa (mean ± SD: 255.4 ± 195.0). Due to the potential for mis-assembly, fragmented assemblies, or mis-annotation in genomes and metagenomes, we removed 2,587 sequences with lengths >2 standard deviations of the mean (636.5 aa), including ~650 proteins >1,000 aa in length. We retained shorter sequences (<2 SD) as they contained key CA domains identified by COG and Pfam IDs. The remaining CA sequences were clustered using MMseqs2 [1] at both 30% and 50% aa similarity. Clustering putative CA sequences at 50% identity doubled the number of clusters compared to 30% identity. However, both methods resulted in a similar number of clustered sequences: 51,391 and 53,289 sequences were sequences at 50% and 30% identity, respectively, after excluding clusters with one (singleton) or two (doubleton) sequences. Similar cluster and sequence counts suggest low sensitivity to identity thresholds. All clusters contained sequences from a single CA class, illustrating no cross-grouping between classes. To avoid mixing distinct functions, downstream analyses used clusters from 50% aa identity.

CA sequences were retrieved from 24,184 genomes or metagenome-assembled genomes (MAGs) representing 85 bacterial, 12 archaeal, 6 algal, and 9 fungal phyla. Information on whether a genome was derived from a cultured or uncultured sample (i.e., MAG) was available only for genomes retrieved from IMG. All fungal genomes in Mycocosm and IMG are from cultured isolates and not metagenomes. Genomes of "algae" in PhycoCosm represent green algae (Chlorophyta), red algae (Rhodophyta), as well as a diverse assemblage of unicellular eukaryotes, including diatoms (Bacillariophyta), amoeboids (Cercozoa), haptophytes (Haptista).

***Taxonomic Assignments of CA Representative Clusters.*** The R packages *taxize* version 0.9.99 [2] and *rentrez* version 1.2.3 [3] were used to retrieve the complete taxonomic classification for each CA sequence based on its NCBI taxonomy identifier (TaxId). Taxonomic information for each sequence was used to determine the lowest common ancestor (LCA) of each cluster (i.e., the lowest taxonomic level shared by all members of the cluster). For example, clusters that contain sequences from closely related taxa are assigned taxonomic ranks near the tips of the NCBI tree, whereas clusters with sequences from more distantly related taxa are assigned to taxonomic ranks closer to the root of the tree (i.e., "higher" taxonomic rank) [4]. We used the Medical Subject Heading [5] term "Gram-Positive Bacteria" to obtain a list of such bacteria and examine their representation in the CA dataset.

***Phylogenetic analyses of CA clades.*** Initial phylogenetic analysis of α-, β-, and γ-CA included 242, 553, and 504 sequences, respectively, representing either reference CA sequences with experimental validation (reference sequences listed in Table S2) or representative sequences from each non-singleton or non-doubleton CA cluster (i.e., cluster representative sequences). We removed representative cluster sequences that did not align to the metal-binding residues [6–9], and/or did not encompass at least 75% of the core CA alignment (i.e., the region of the MSA containing >95% of the sequences). The removal of 257 CA cluster sequences (220 representing γ-CA) substantially decreased the number of γ-CA sequences included in phylogenetic analysis (i.e., from 504 to 284 γ-CA sequences). Phylogenetic trees from IQ-TRee were visualized with the Interactive Tree Of Life (iTOL) version 5 [10], Evolview version v3 [11], and T-BAS version 2.1 [12].

### ***<u>RESULTS</u>***

***LCA taxonomy across the CA phylogenies.*** In addition to fungal-specific β-D subclade (β-D3), β-D1 subclade contained only CA from Bacteria and Archaea, whereas β-D2 contained a greater taxonomic diversity of CA (Fig. 4a). Multiple β-A subclades appear specific to either Basidiomycota (β-A2, A7), Ascomycota (β-A3, A4, A5), or non-dikaryotic fungi (β-A3), which were interspersed with bacterial and algal CA in subclade (β-A6). Fungal CA was also found in subclade β-A1 with CA from Bacteria, Algae, and other Eukaryotes. The β-E clade is comprised of CA with LCAs assigned only to Archaea and Bacteria, similar to its sister β-C clade (Fig. 4a).

The β-B clade contained three major subclades of CA that were assigned to both eukaryotes and bacteria. Represented by numerous reference CA, the γ-A1 clade is comprised of CA clusters assigned to Bacteria, Algae, and other Eukaryotes, as well as CA clusters shared across multiple Superkingdoms (Fig. 4c). In contrast, the γ-B clade is comprised of CA primarily from Bacteria (including numerous reference sequences) and Archaea. The γ-A2 clade contains CA assigned primarily to Algae and other Eukaryota. The largely unresolved γ-A3 clade contains the majority of γ-CA clusters representing CA assigned to all Superkingdoms, including subclades that appear specific to Fungi and other eukaryotes (Fig. 4c). For example, the α-A and α-B clades were comprised of CA clusters with the lowest common ancestor (LCA) assigned to Bacteria, Algae, and Eukaryota (Fig. 4b). The α-C clade contains two subclades: one clade has CA assigned only to Fungi and another has CAs assigned to both Eukarya and Bacteria (Fig. 4b). The α-D clade also is comprised of CA assigned only to Fungi, whereas the α-E was primarily comprised of bacterial CA (Fig. 4b).

***Distribution of CA classes across metaT/G datasets.*** Numerous CAs were observed in studies of engineered ecosystems derived from cities; however, environmental samples derived from the air registered the fewest number of CA hits (n = 91) (Fig. 5a, Table S8, Additional file 5). On average, α- and β-CA clusters had hits in <5% of the datasets (mean = 1.9% and 4.9%, respectively), whereas on average γ-CAs occurred in 13% of datasets (P<0.0001). The majority of CA were infrequently observed across the metagenomic/metatranscriptomic datasets (i.e., <20%); however, a small fraction of CA clusters were observed in >50% of datasets (Fig. S11a). In addition, the frequency distribution of CA classes differed as a function of LCA assignment (Fig. S11b). The mean frequency and mean number of hits for CA clusters in the metaG/T datasets also differed as a function of LCA taxonomic assignment for each CA class (Fig. S12).

***Abundance of γ-CA in metaG/T datasets.*** Our analysis revealed an unexpected abundance of γ-CA across all environments compared to previous results from Meredith et al. [13]. One potential explanation for this discrepancy is the cluster size threshold (i.e., here, we used a minimum of three sequences per cluster, whereas previous analyses used a minimum of 50 sequences [13]). To assess the impact of cluster size threshold on our results, we examined whether the number of clusters with few sequences (i.e., <4) differed between CA classes. We observed similar numbers of smaller-sized clusters for both β- and γ-CA (89 β-CA clusters and 105 γ-CA clusters with <4 sequences), and for both classes, smaller clusters comprised 19% of the total number of clusters/class. We also found a similar number of clusters with >49 sequences for each class (62 β-CA and 57 γ-CA). These results suggest that the abundance of hits to γ-CA HMMs in metaG/T datasets is unrelated to the CA cluster size threshold used.

***Potential bias in metaT/G results.*** One plausible explanation for the over-abundance of HMM hits to bacterial CA is the strong bias in metaG/T sequences toward bacterial reads [14]. In addition, some environments were sampled more frequently than others in the metaG/T database (Table S8). For example, there were only five hits to host-associated environments in metaT datasets compared to >78k hits to metaG datasets for that same environment, which reflects the few metaT datasets from eukaryotic hosts, as these typically only result in host reads. To address these biases, we performed analyses on all HMMs, as well as on only HMMs with the highest number of hits to the metaG/metaT database at IMG per ecosystem category (i.e., the top 10 or 50 most abundant CA). All analyses supported the wide distribution of CA from α-, β-, and γ-classes across most sampled ecosystems, including both aquatic and terrestrial environments (Fig. 6, Fig. S15, Fig. S16). To account for the over-abundance of bacterial/archaeal reads in metaG/T datasets, we also analyzed profiles of prokaryotic and eukaryotic CA separately. In doing so, we found that sampling the top ten and fifty HMMs per ecosystem reproduced the same results for fungal α-CAs and did not substantially differ for fungal β-CAs. However, analysis of the top 50 CA substantially increased the number of HMMs present in all ecosystems except for air (Fig. S14, Table S9).

### ***LIST OF SUPPLEMENTARY TABLES***

**Table S1.** CA protein families, their description, and number of CA recovered from IMG/MER, MycoCosm, and PhycoCosm (a), IMG/MER (b), MycoCosm (c), or PhycoCosm (d). Different approaches were used, such as BLAST searches or sequence retrieval using COG, Pfam, KEGG, Interpro, and EC IDs. The values are based on the published genomes as of May 5th, 2021.

**Table S2.** Reference sequences used in the phylogenetic analyses.

**Table S3.** Table showing the number of CA sequences per database and group of organisms (a), the number of genomes containing at least one CA per database and group of organisms (b), the number of genomes without CA per database and group of organisms (c), and the number of genomes with a single class of CA, two classes of CA, and the three classes of CA. The values are based on the published genomes as of May 5th, 2021.

**Table S4.** Fungal and algal genomes without detectable CA genes.

**Table S5.** Frequency of CA per genome (a), also detailed by database (b), superkingdom (c), and group of organisms (d). The values are based on the published genomes as of May 5th, 2021.

**Table S6.** Clustering summary, after clustering sequences at 50% (a, b) and 30% (a, c) sequence identity thresholds.

**Table S7.** Percentage of α-, β-, and/or γ-CA clusters unique to a specific superkingdom (a), kingdom (b), phylum (c), class (d), order (e), family (f), and genus (g).

**Table S8.** Count of HMM hits to the metaG/metaT database at IMG per environment. (a, b) α-CA HMMs. (c, d) β-CA HMMs. (e, f) γ-CA HMMs.

**Table S9.** Ten and 50 most retrieved HMMs per ecosystem, after subsampling archaeal and bacterial HMMs (a-f) or fungal HMMs (g-l). HMMs were queried against the metaG/metaT database at IMG.

### ***SUPPLEMENTARY FIGURES***

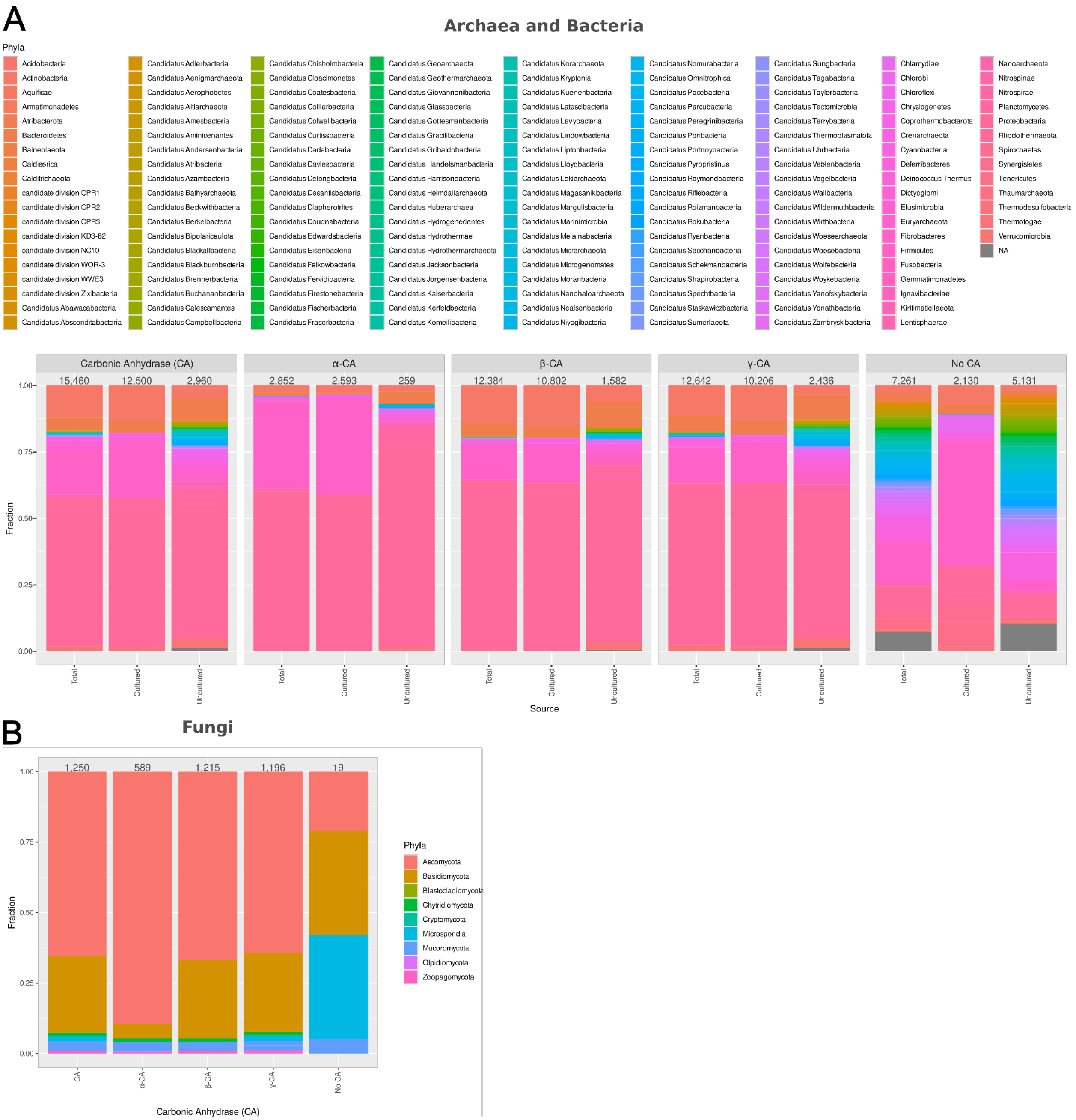


**Figure S1.** Relative abundance of (A) Bacteria and Archaeal and (B) Fungal Phyla with and without CA derived from genomes of cultured and uncultured organisms. The number at the top of each bar indicates the number of genomes in the corresponding category. Also see Figure 1.

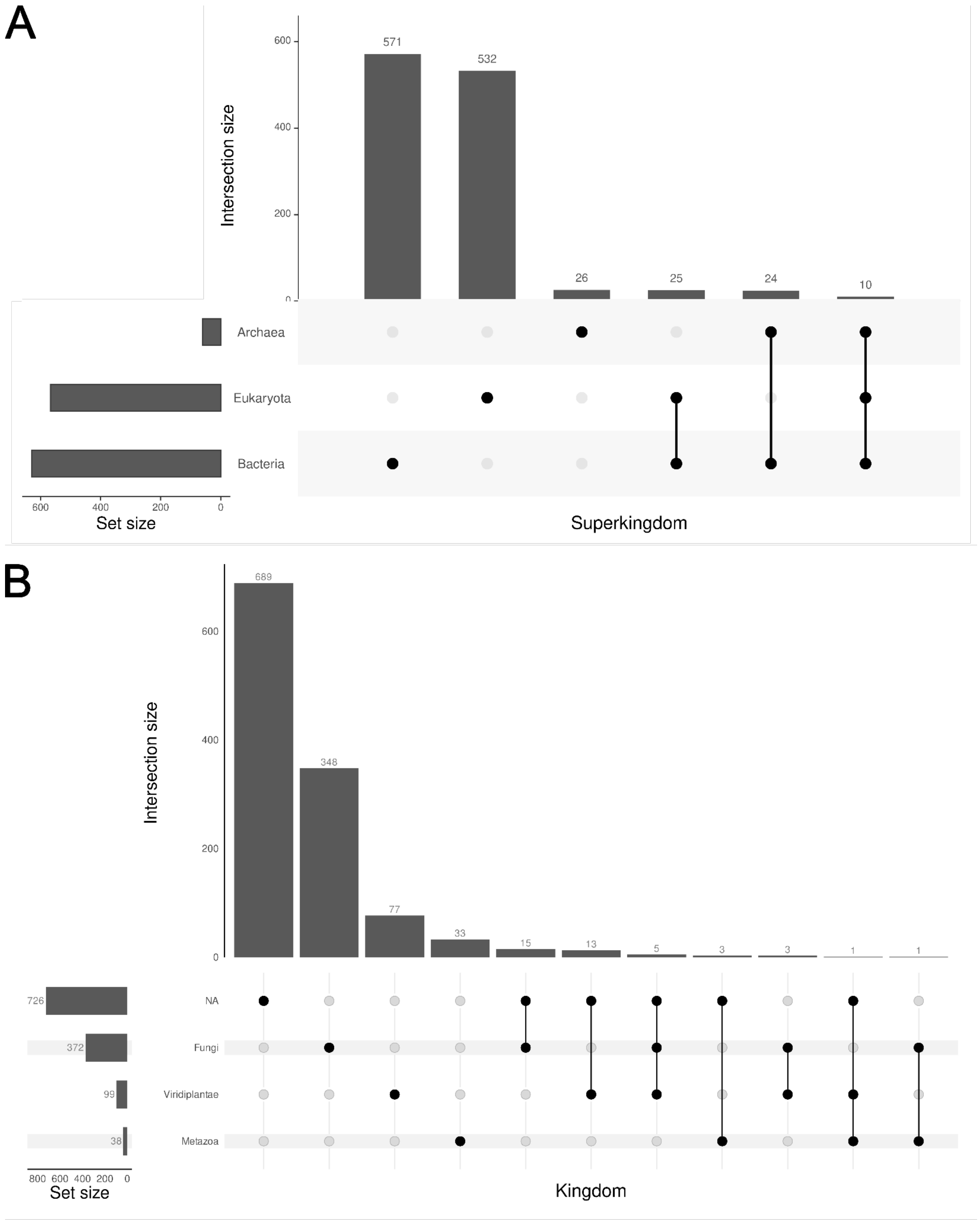


**Figure S2.** (**A**) UpSet plot displaying the intersection of different (A) Superkingdoms and (**B**) Kingdoms for clusters with more than two sequences.

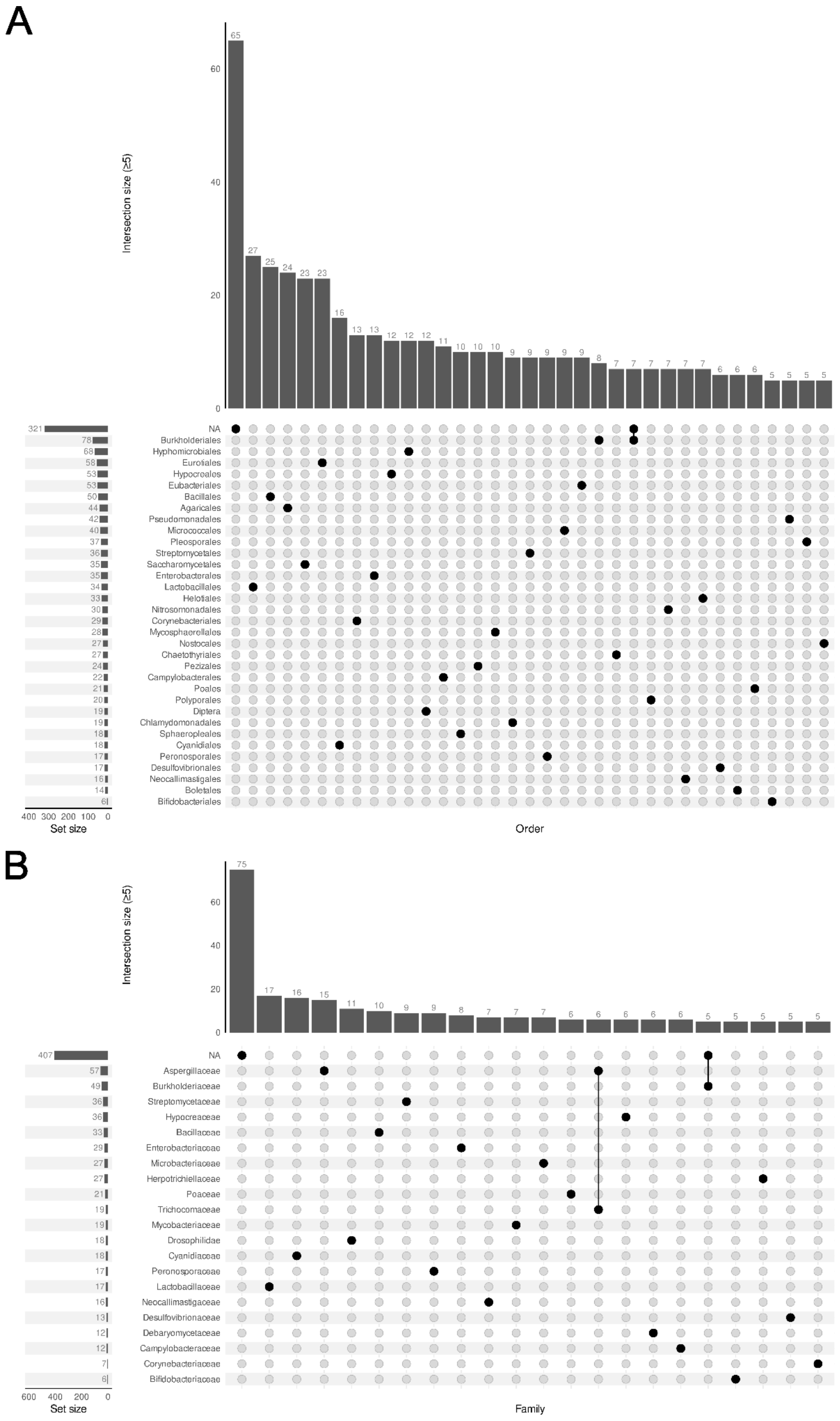


**Figure S3.** UpSet plot displaying the intersection of different (A) orders and (B) families for clusters with more than two sequences. Only intersection sizes greater than or equal to five are shown.

α-CA

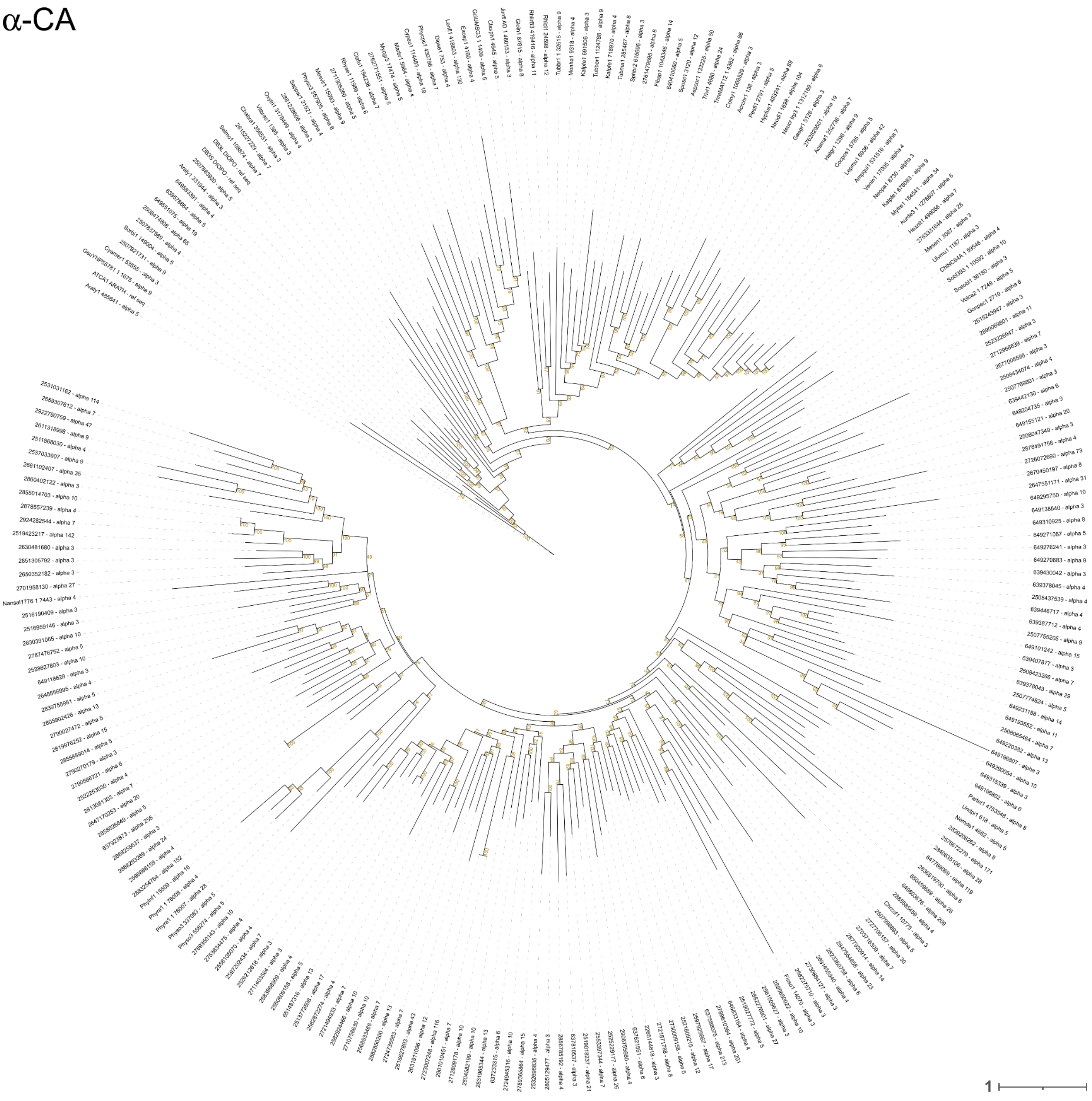

**Figure S4.** Phylogenetic tree of α-CAs. The phylogeny was reconstructed using the cluster representative sequences and the reference sequences by a maximum-likelihood approach with IQ-TREE multicore version 1.6.12, employing the best-fitting model of sequence evolution selected by ModelFinder. Node support was evaluated using 1000 ultrafast bootstrap (UFBoot) replicates. Node support is indicated in mustard. The scale bar represents the number of substitutions per site.

β-CA

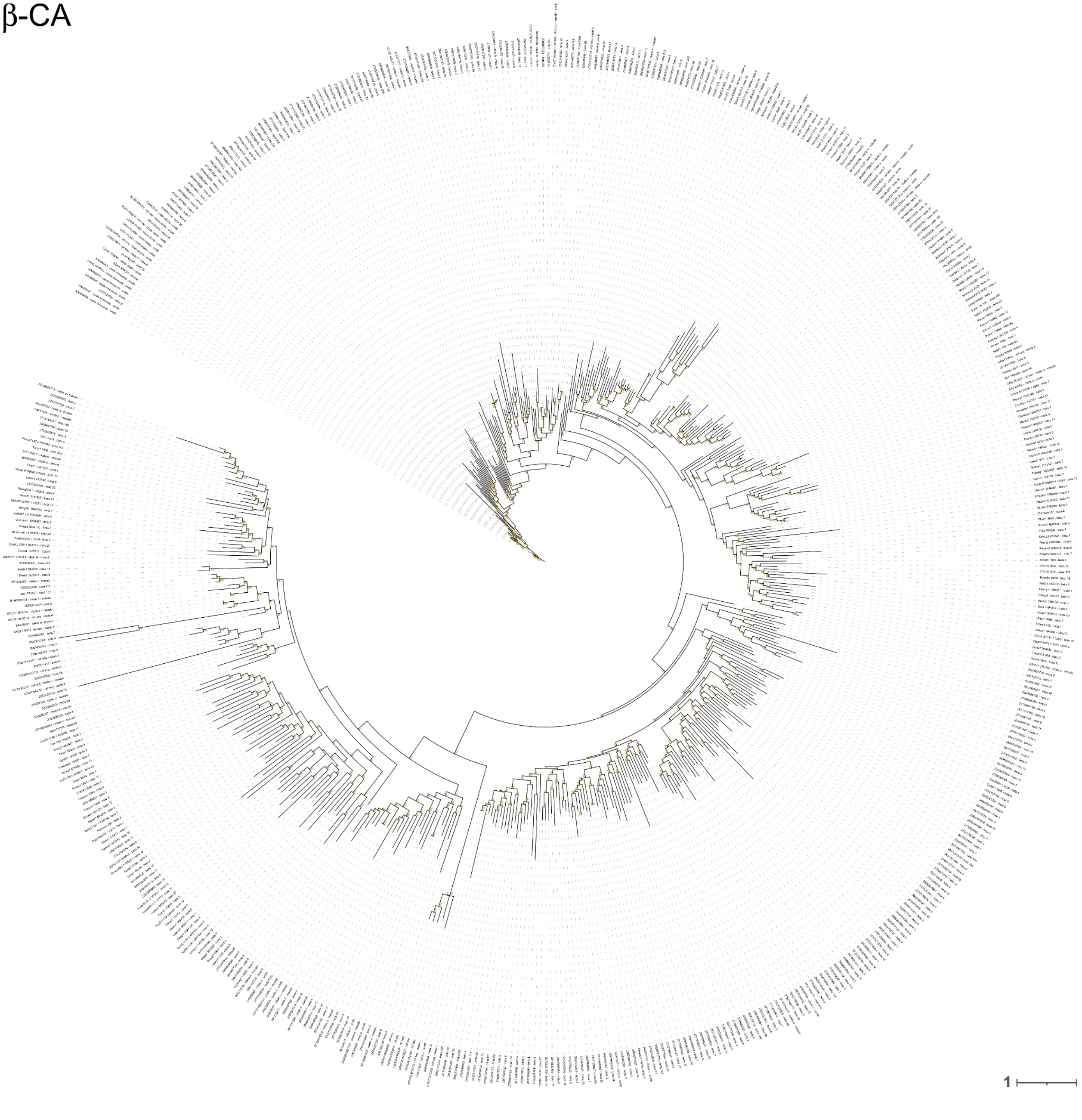


**Figure S5.** Phylogenetic tree of β-CAs. The phylogeny was reconstructed using the cluster representative sequences and the reference sequences by a maximum-likelihood approach with IQ-TREE multicore version 1.6.12, employing the best-fitting model of sequence evolution selected by ModelFinder. Node support was evaluated using 1000 ultrafast bootstrap (UFBoot) replicates. Node support is indicated in mustard. The scale bar represents the number of substitutions per site.

γ-CA

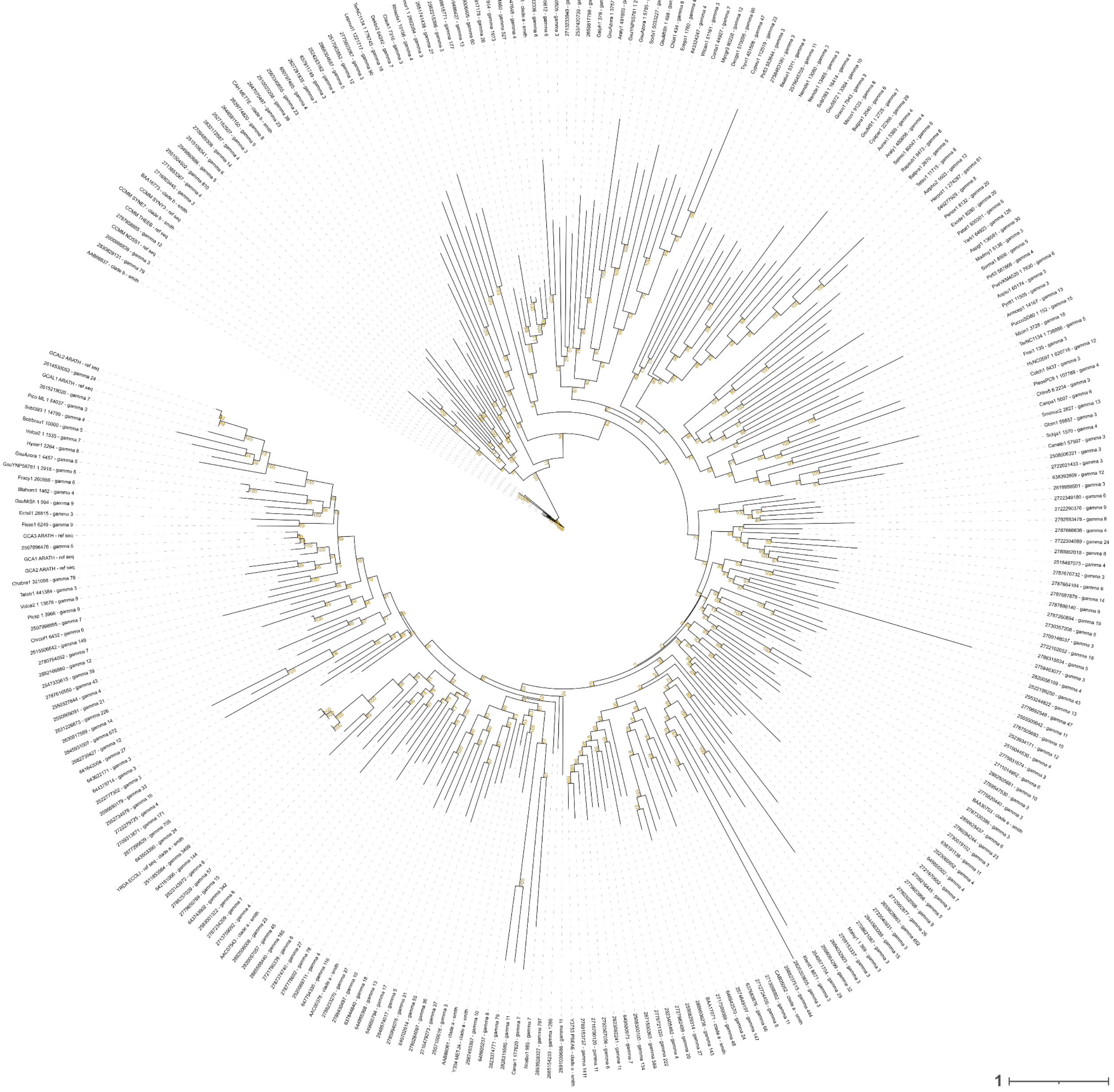

**Figure S6.** Phylogenetic tree of γ-CAs. The phylogeny was reconstructed using the cluster representative sequences and the reference sequences by a maximum-likelihood approach with IQ-TREE multicore version 1.6.12, employing the best-fitting model of sequence evolution selected by ModelFinder. Node support was evaluated using 1000 ultrafast bootstrap (UFBoot) replicates. Node support is indicated in mustard. The scale bar represents the number of substitutions per site.

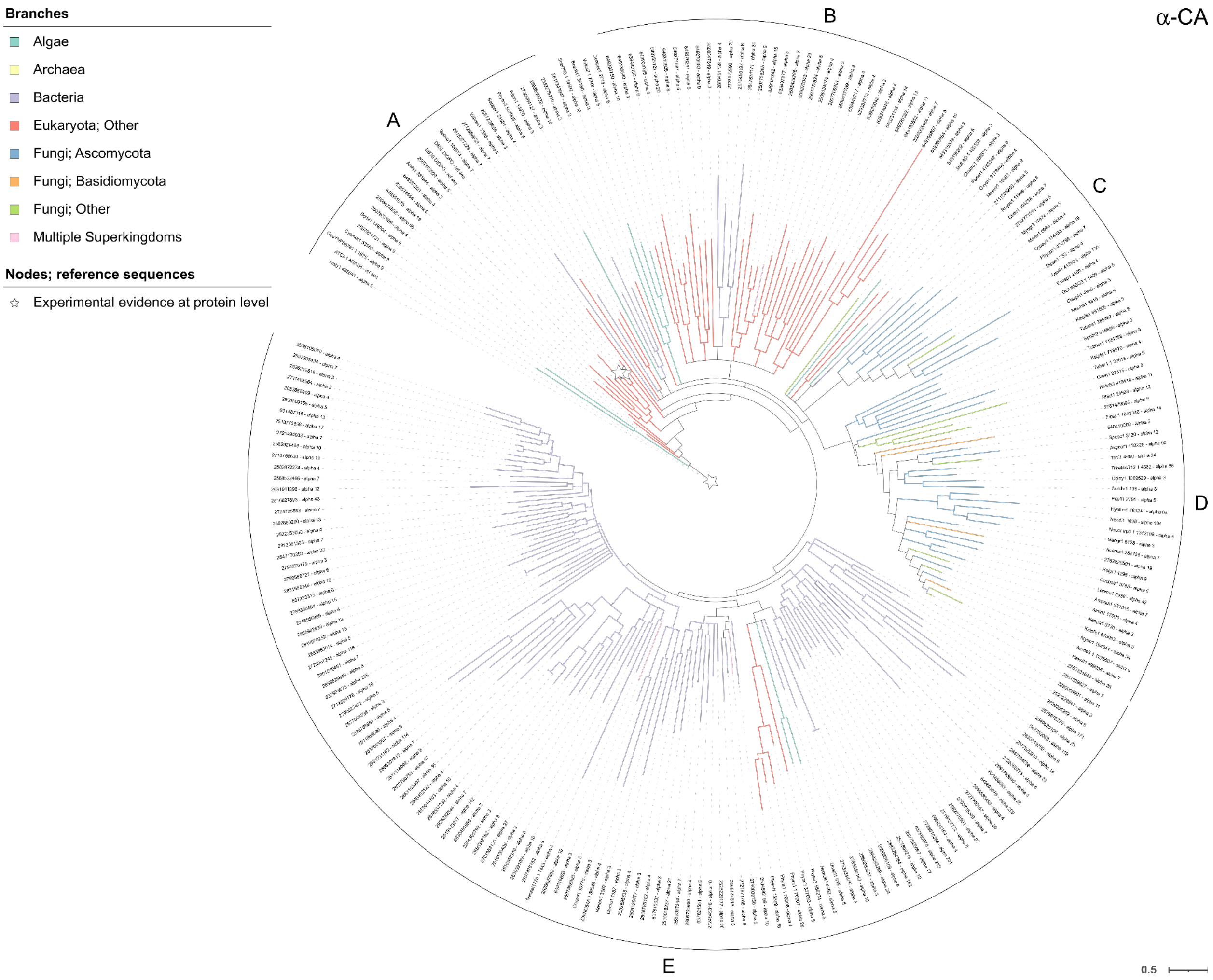


**Figure S7.** Phylogeny of α-CA after collapsing nodes with low support values. The phylogeny was reconstructed using the cluster representative sequences and reference sequences by a maximum-likelihood approach with IQ-TREE multicore version 1.6.12 employing the best-fitting model of sequence evolution selected by ModelFinder. Node support was evaluated using 1000 ultrafast bootstrap (UFBoot) replicates. Nodes with less than 95% UFBoot support were collapsed. The scale bar represents the number of substitutions per site. Branches are colored according to the cluster representative taxonomy assigned by the lowest common ancestor-like approach. The stars indicate reference sequences labeled as 'Experimental evidence at protein level' in UniProtKB.

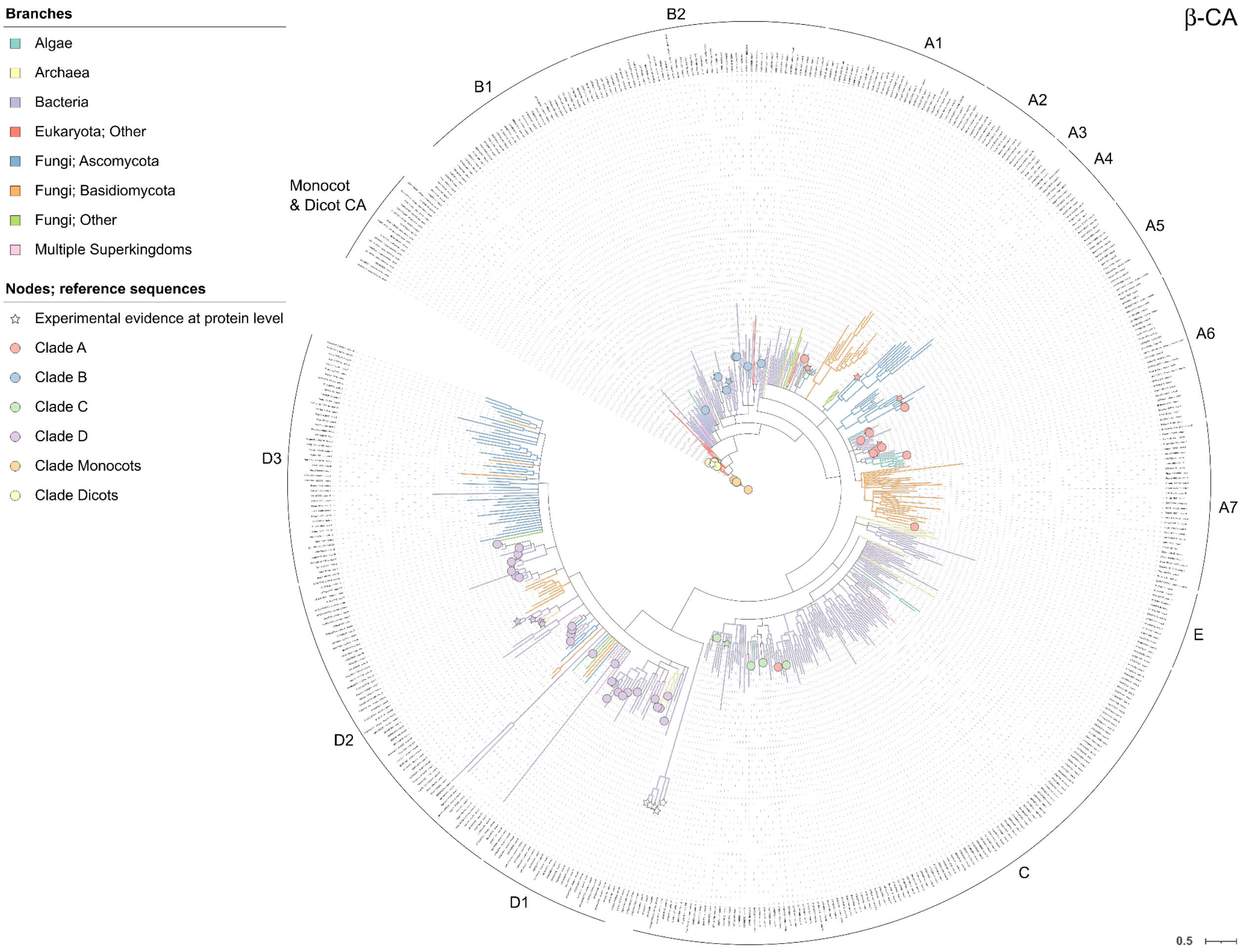


**Figure S8.** Phylogeny of β-CA after collapsing nodes with low support values. The phylogeny was reconstructed using the cluster representative sequences and reference sequences by a maximum-likelihood approach with IQ-TREE multicore version 1.6.12 employing the best-fitting model of sequence evolution selected by ModelFinder. Node support was evaluated using 1000 ultrafast bootstrap (UFBoot) replicates. Nodes with less than 95% UFBoot support were collapsed. The scale bar represents the number of substitutions per site. Branches are colored according to the cluster representative taxonomy assigned by the lowest common ancestor-like approach. Reference sequences are indicated with a symbol at the edge of the tip, which is colored based on the clade assignment Smith et al. (1999) and Masaki et al. (2021). The stars indicate reference sequences labeled as 'Experimental evidence at protein level' in UniProtKB.

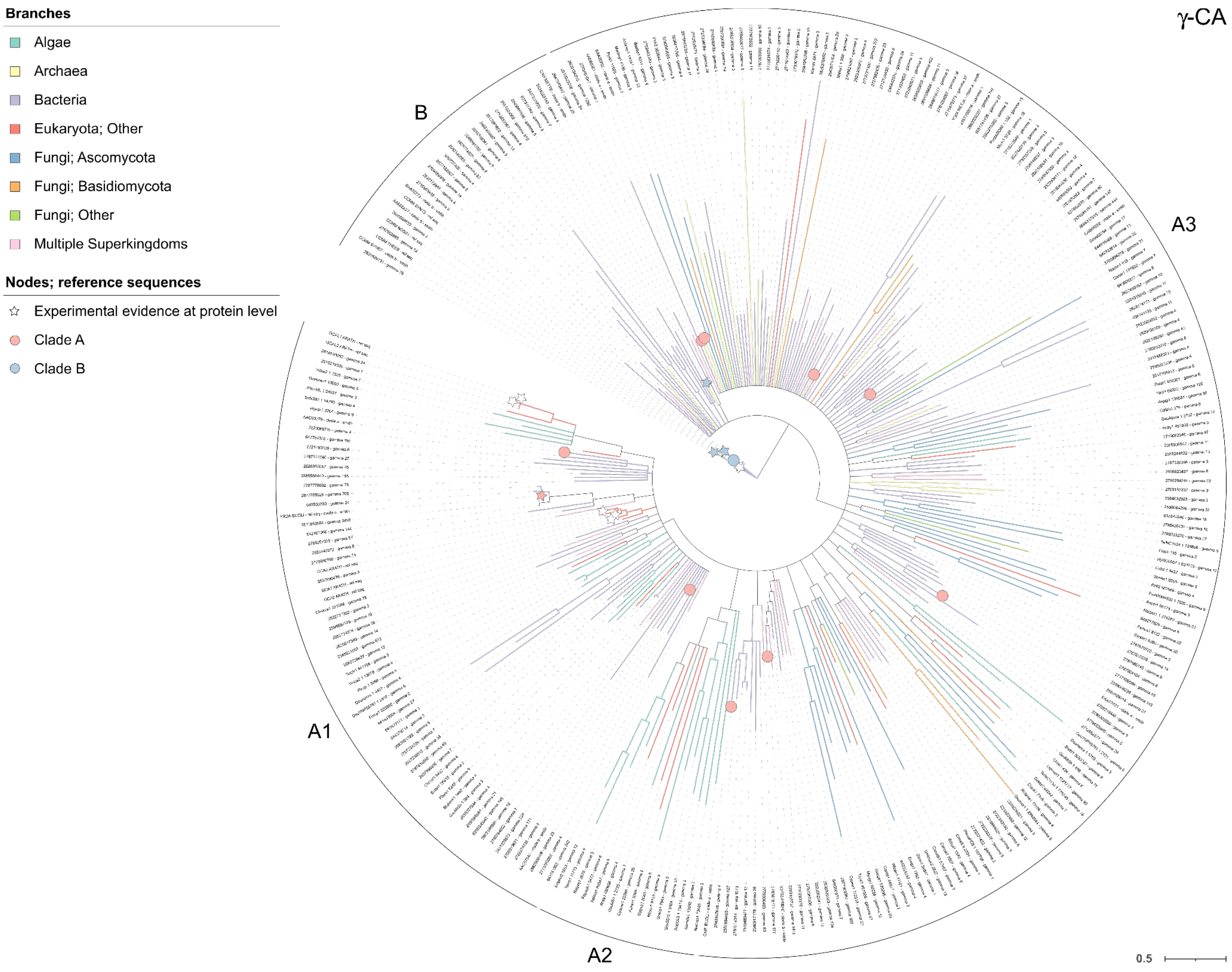


**Figure S9.** Phylogeny of γ-CA after collapsing nodes with low support values. The phylogeny was reconstructed using the cluster representative sequences and reference sequences by a maximum-likelihood approach with IQ-TREE multicore version 1.6.12 employing the best-fitting model of sequence evolution selected by ModelFinder. Node support was evaluated using 1000 ultrafast bootstrap (UFBoot) replicates. Nodes with less than 95% UFBoot support were collapsed. The scale bar represents the number of substitutions per site. Branches are colored according to the cluster representative taxonomy assigned by the lowest common ancestor-like approach. Reference sequences are indicated with a symbol at the edge of the tip, which is colored based on the clade assignment Smith et al. (1999). The stars indicate reference sequences labeled as 'Experimental evidence at protein level' in UniProtKB.

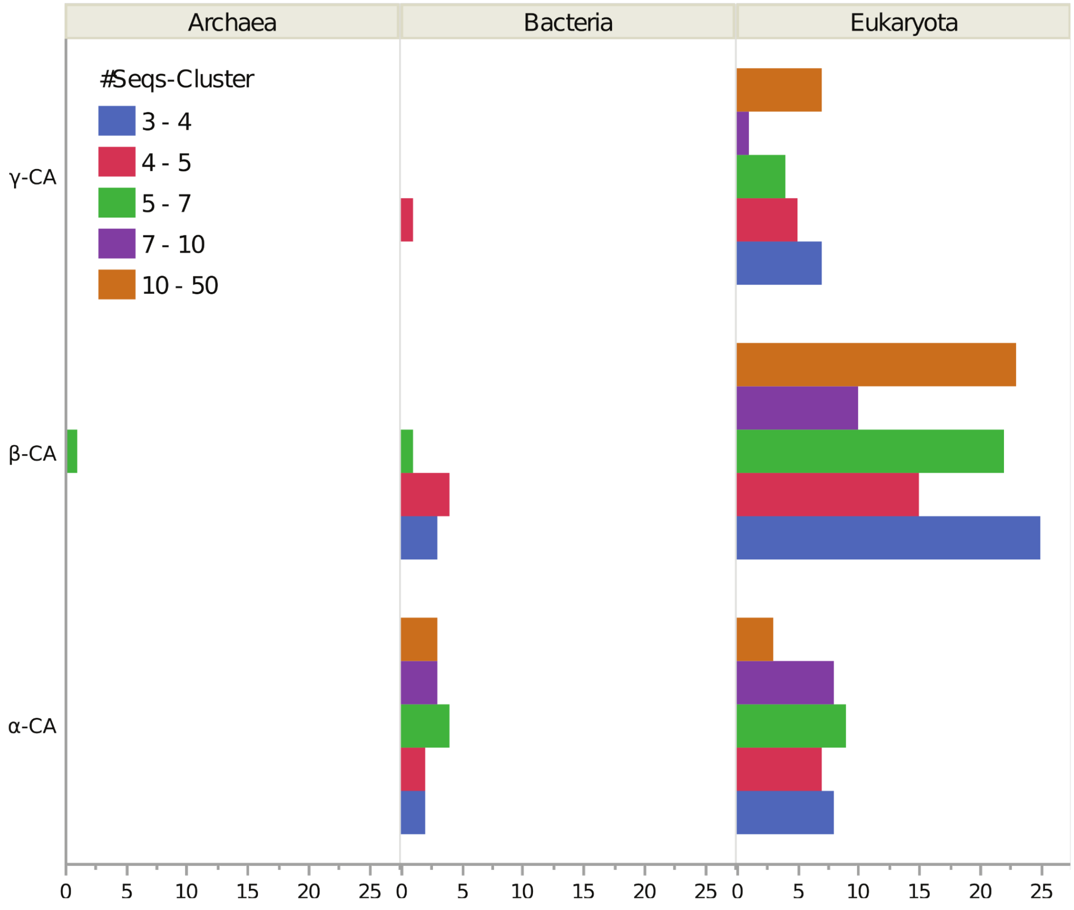


**Figure S10.** Histogram of cluster size for α-, β-, γ-CA clusters with HMM that did not match any of the CA sequences obtained from previous metagenomic and metatranscriptomic studies. Bars are colored by the number of sequences in each CA cluster.

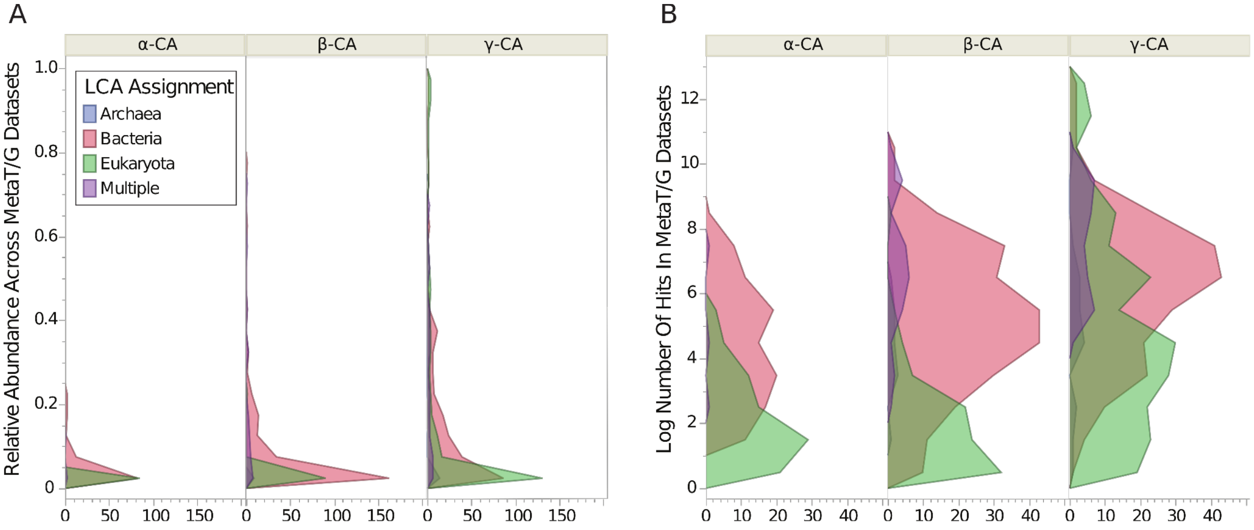


**Figure S11.** Frequency distribution of the (**A**) relative abundance of each CA class out of 3,672 metagenomic/metatranscriptomic datasets and (**B**) log number of total hits for each CA class. Colors indicate the LCA taxonomic assignment of each CA cluster.

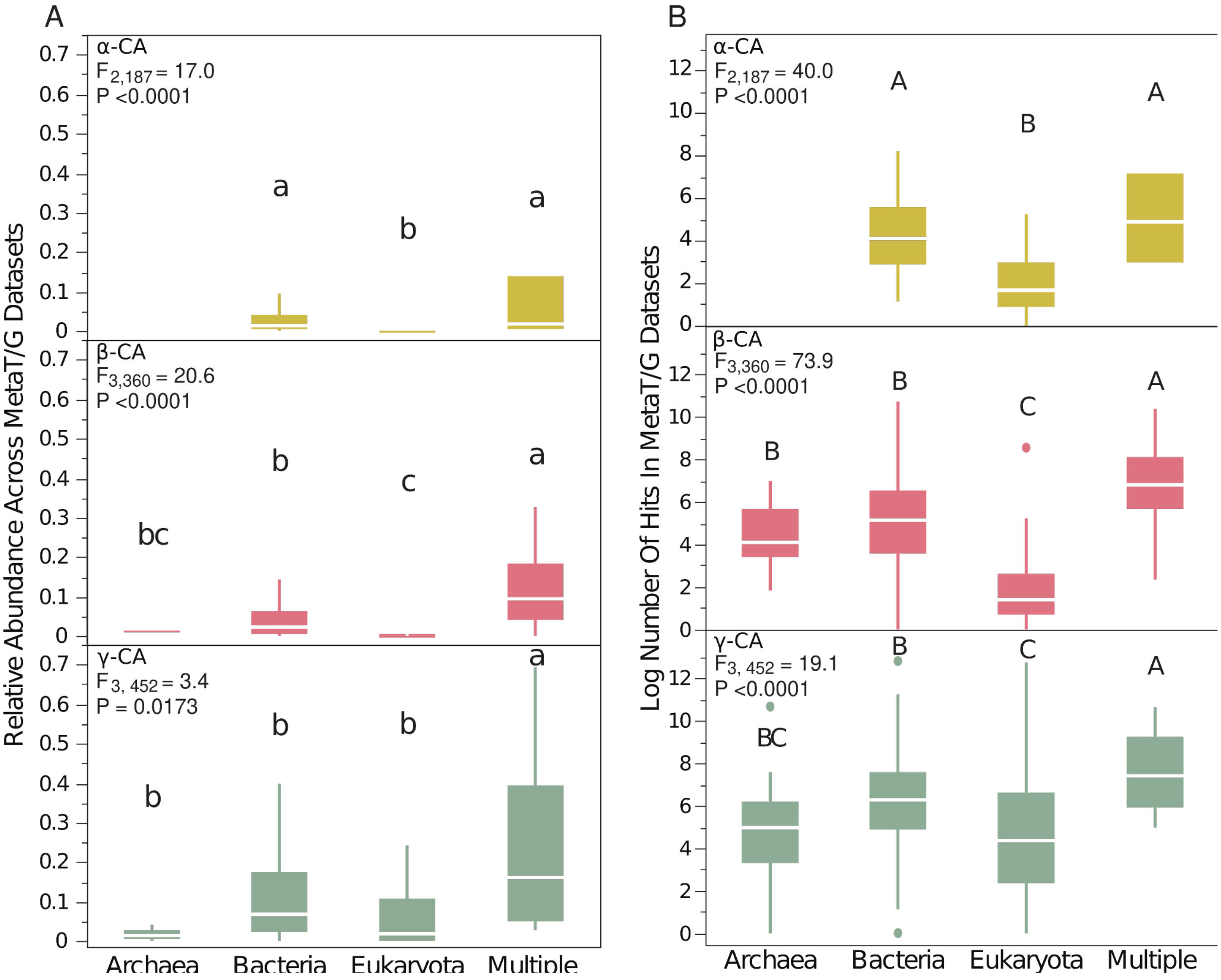


**Figure S12.** Median (**A**) relative abundance of α-, β-, and γ-CA clusters out of 3,672 metagenomic/metatranscriptomic datasets and (**B**) log number of total hits for each CA class as a function of the LCA taxonomic assignment of each CA. "Multiple" indicates CA clusters comprised of sequences from multiple Superkingdoms. Statistical results from one-way ANOVA for each CA class are shown in the upper left of each panel; letters correspond to differences after post-hoc Tukey's HSD ($P<0.05$). Quantile box plots show the minimum and maximum values (vertical lines), the 50th percentile (i.e., median; white line), and the 25th to 75th percentiles (area of the box).

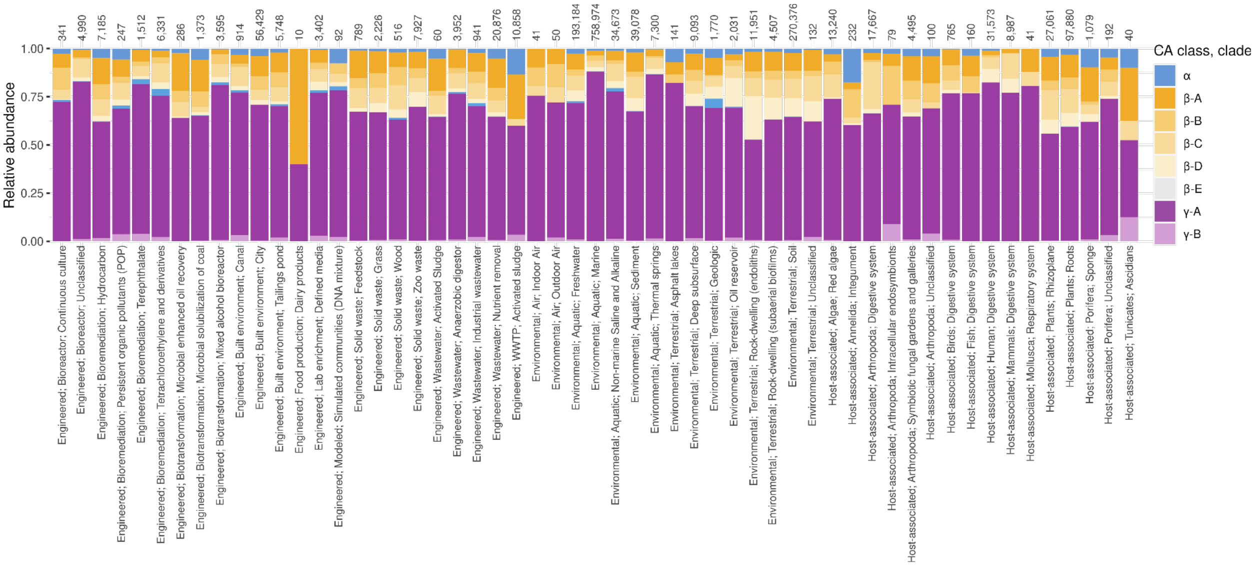


**Figure S13. Relative abundance of CA clades and classes by all environmental categories**. Stacked bar chart showing the relative abundance of α-, β- and γ-CA HMM hits to the combined metagenomic/metatranscriptomic database downloaded from IMG. Numbers on the top of each bar represent the total number of hits per environment. CA class. CA clade information follows Fig. 4. Although the greatest fraction of hits for each environment was to γ-CA, among the β-CAs, β-D CAs appear relatively abundant in terrestrial environments, particularly in rock-dwelling (endoliths) and soils.

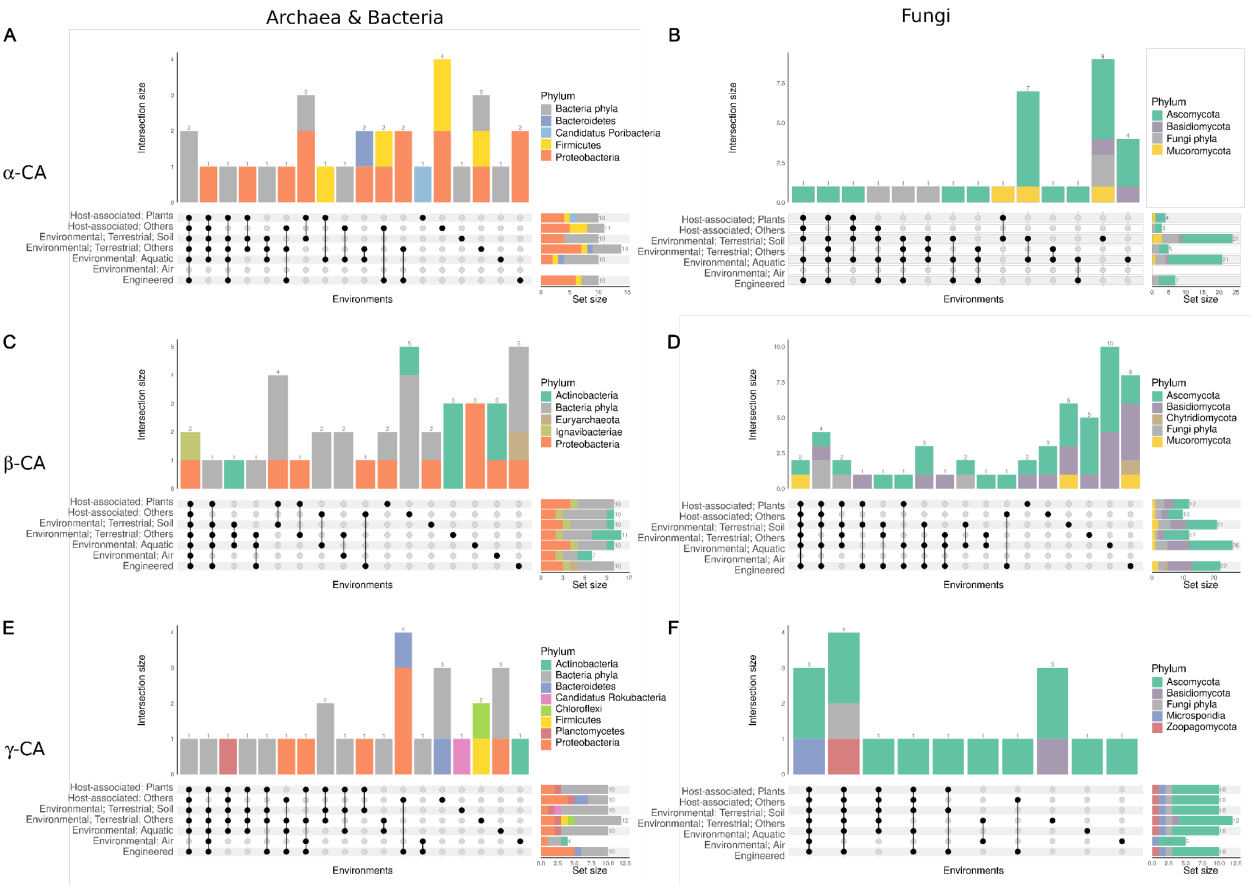


**Figure S14.** UpSet plot displaying the intersection of different ecosystems for the 10 most retrieved HMMs per ecosystem. Archaeal and Bacterial HMMs (a, c, e), and Fungal HMMs (b, d, f) were subsampled and analyzed separately. Bars are colored by phyla, which was assigned by the LCA approach. See Fig. 6 for the contributions of all CA classes together. The most abundant HMMs are listed in Table S8.

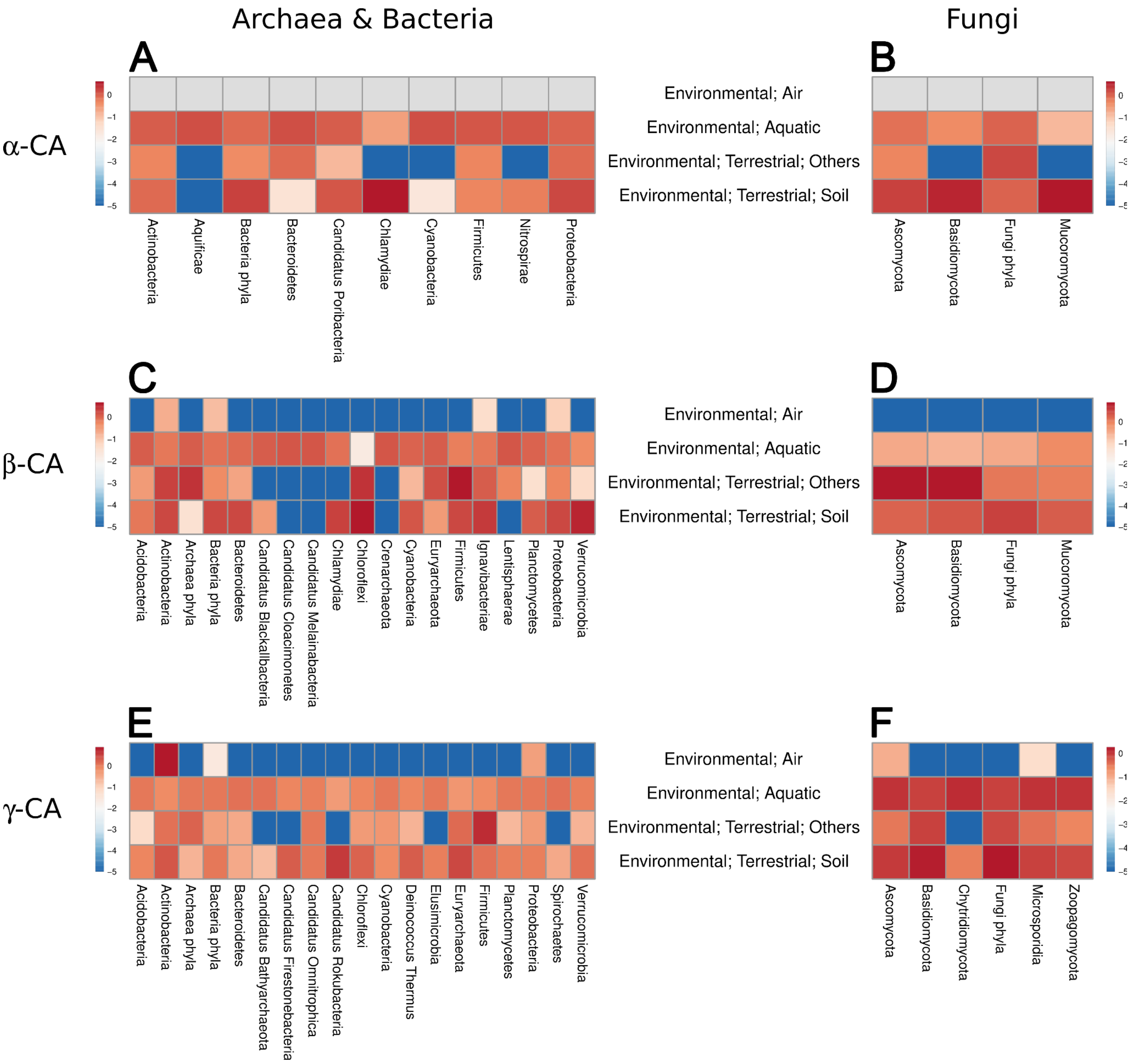


**Figure S15.** Heatmap showing the relative abundance of α-, β- and γ-CA HMM hits to the metaG/metaT database at IMG from different archaeal and bacteria or fungal phyla across the major environmental ecosystems for all HMMs. Archaeal and bacterial HMMs (a, c, e), and fungal HMMs (b, d, f) were subsampled and analyzed separately. The number of hits was scaled by the proportion of studies for the respective ecosystem and the abundance of each phylum and log-transformed. The phyla were assigned by the LCA-like approach.

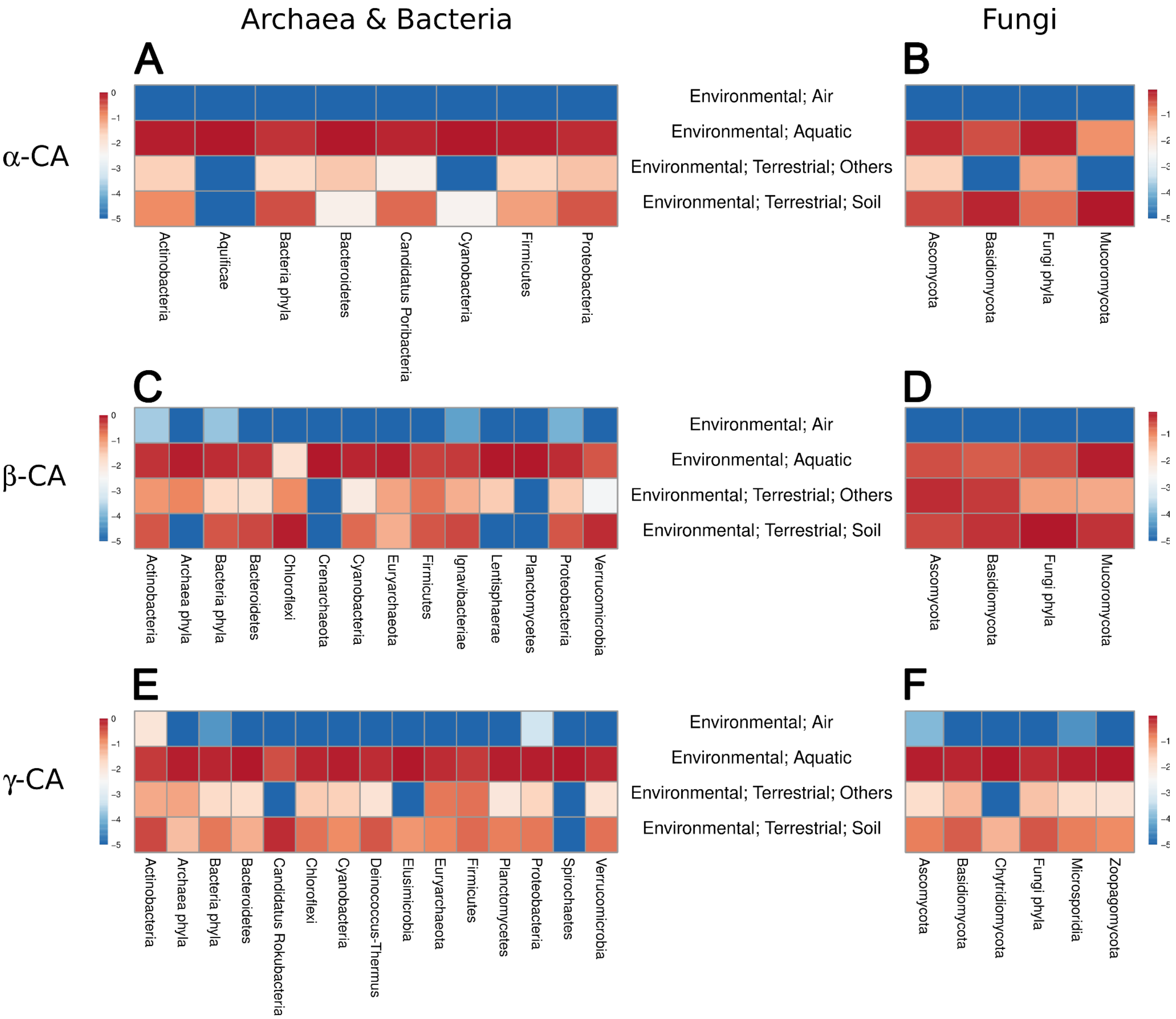


**Figure S16.** Heatmap showing the relative abundance of α-, β- and γ-CA HMM hits to the metaG/metaT database at IMG from different archaeal and bacteria or fungal phyla across the major environmental ecosystems for the ten most retrieved HMMs per ecosystem. Archaeal and bacterial HMMs (a, c, e), and fungal HMMs (b, d, f) were subsampled and analyzed separately. The number of hits was scaled by the abundance of each phylum, and log-transformed. The phylum was assigned by the LCA-like approach.

*ADDITIONAL FILES (available at Figshare: 10.6084/m9.figshare.28156742)*

**Additional file 1.** Tab-separated file providing the identification, source, and taxonomy for the genes included in this study.

**Additional file 2.** Cluster FASTA-like format produced by MMseqs2 after clustering the sequences at 50% sequence similarity. Two successive identical names indicate a new cluster; the first sequence is the cluster representative sequence, which also gives the group its name.

**Additional file 3.** Tab-separated values (TSV) file showing the lowest taxonomic level shared by all the members in the cluster. The taxonomic ranks analyzed were superkingdom, kingdom, phylum, clade, class, order, family, genus, species, and strain.

**Additional File 4.** Edited multiple sequence alignments and corresponding phylogenetic trees for α-, β-, and γ-carbonic anhydrases (CAs).

**Additional file 5.** Count of CA HMM hits to the metaG/metaT database at IMG. GOLD Ecosystem classification is provided.
